# A single-cell atlas of *Toxoplasma* sexual development in the feline intestinal tract

**DOI:** 10.1101/2025.10.31.685900

**Authors:** Hisham S Alrubaye, Sarah M Reilly, Rafaela da Silva, NyJaee Washington, Jon P Boyle

## Abstract

*Toxoplasma gondii* undergoes sexual development exclusively in the feline intestine, a process critical for genetic diversity and population expansion. Recent studies have identified genes critical in suppressing presexual development and metabolic differences in felines that may promote sexual development, but to date the gene regulatory networks driving development in the cat are unknown. To investigate this, we performed single-cell transcriptomics on parasites isolated from cat intestines, using fluorescent reporter strains and flow cytometry. From 15,068 cells across two experiments, we identified rare populations, including cells that bear all of the hallmarks of gametes. Candidate genes emerging from this study were tested via CRISPR-Cas9 Perturb-seq, identifying AP2X6 as a regulator of macrogametocyte development. Our single-cell data extends what is known about gene expression changes throughout sexual development and should be useful to those in the field working towards inducing gametogenesis, mating, and oocyst production *in vitro*.

## Main

*T. gondii* has a complex life cycle that includes both asexual and sexual reproduction. While asexual stages can develop in nearly all cells of warm-blooded animals, sexual development is restricted to the epithelial cells of the felid small intestine. The sexual phase of *T. gondii* is essential for generating genetic diversity through recombination and for large scale environmental dissemination. A single feline host is capable of shedding almost a billion oocysts after a single infection ^1–4^. Shed oocysts mature further after completion of meiosis and one round of mitosis, resulting in the production of infectious sporozoites.

Among the developmental stages of *T. gondii*, the tachyzoite and bradyzoite have been studied the most due to their relative tractability *in vitro*, and a substantial amount of information is available on the mechanisms of tachyzoite to bradyzoite stage transition^5,6^ as well as stage-specific cell cycle regulation^7,8^, metabolism^9,10^, and host-parasite interactions^11^. In contrast, due to challenges associated with cultivating pre-sexual and ultimately sexual stages, there is far less known about how this developmental program is triggered and executed by the parasite during infection of the feline host. Early foundational ultrastructural studies in the 1970s began to differentiate between the asexual and sexual enteric stages of *T. gondii*^12,13^. These and other studies identified five morphologically distinct pre-gamete stages that develop following infection of cat enterocytes^14^, providing a developmental framework but the molecular mechanisms underlying these processes remained unknown until recent advances.

Transcriptomic studies showed that *T. gondii* undergoing enteric development in the feline host have unique transcriptional profiles compared to other developmental stages. These bulk transcriptomic studies have been instrumental in uncovering significant transcriptional differences between developmental stages in intermediate and definitive hosts^15–17^. Although reproducibility has been limited, metabolic cues from the feline host have been proposed to trigger the transition from asexual to presexual development in *T. gondii*, suggesting that host-specific signals may contribute to parasite differentiation^18^. In addition, recent studies have identified parasite transcription factors AP2XII-1 and AP2XI-2 as key suppressors of *T. gondii* pre-sexual development, and their genetic depletion *in vitro* permits presexual development^19^. These studies have paved the way for targeted investigations into the regulation of this developmental program^19^. However, the depletion of those suppressors did not yield gametes *in vitro*, indicating that additional factors are required for gametogenesis.

Here, we interrogated the transcriptional program underlying the enteric stages of *T. gondii* in the cat using single-cell RNA sequencing (scRNA seq) to identify previously unknown regulators of enteric development. We identified multiple transcriptionally distinct clusters within the enteric stages, including putative male and female gametocyte populations based on sex-specific expression signatures. To functionally characterize key regulators of parasite development, we designed a targeted perturb-seq screen of 28 candidate genes, leading to the identification of AP2X6 as a regulator of macrogametocyte differentiation. Our data provides the field with novel gene markers for studying distinct subpopulations of *T. gondii*, offering new targets for individual investigation to enhance our understanding of the molecular mechanisms underlying the parasite development in the definitive host.

## Results

### Generation of a cat-competent *Toxoplasma gondii* reporter strain

Previous transcriptomic studies of feline enteric stages relied on bulk RNA-seq, which lacks resolution to capture heterogeneity and rare subpopulations such as transitional forms and gametes^16,17^. To address this limitation, we generated single-cell transcriptomic profiles of parasites isolated from the feline small intestine. We engineered a TgVEG reporter line expressing GFP from a tachyzoite promoter (GRA1) and mCherry from a hybrid promoter (GRA11A + GRA1 upstream sequences) that resulted in a strain expressing GFP during asexual development and mCherry during both asexual and sexual development (Fig. 1A; Extended Data Fig. 1).

**Fig. 1:**
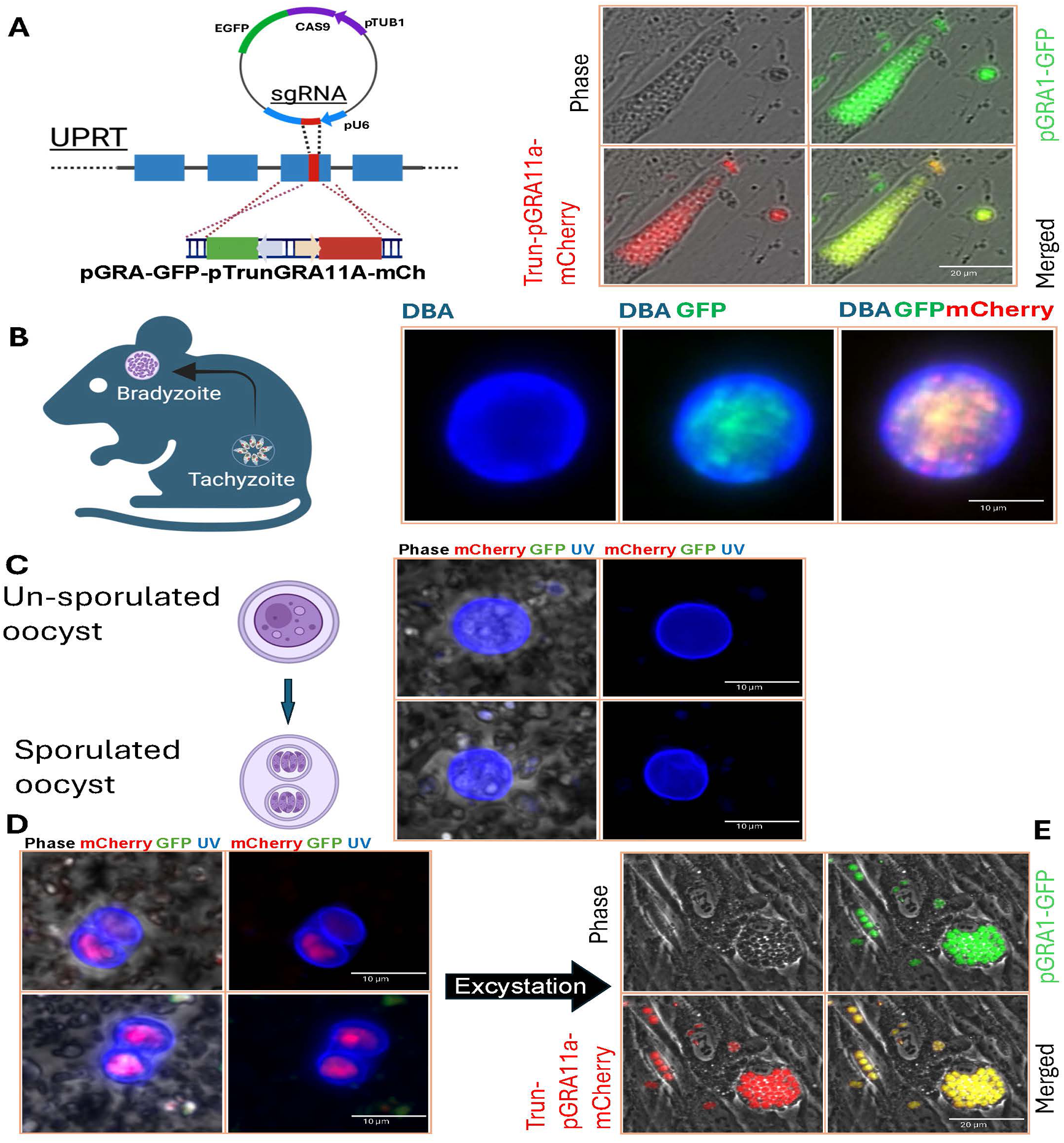
Generation of a cat-competent *Toxoplasma gondii* reporter strain. **A)** A cat-competent (i.e., oocyst-forming) strain, TgVEG, was used in this study. A dual-reporter cassette was constructed with GFP driven by the *GRA1* promoter and mCherry under the control of a truncated *GRA11A* promoter (left panel). TgVEG harboring this reporter construct expressed both GFP and mCherry under tachyzoite conditions (right panels). **B)** Tissue cysts derived from the TgVEG reporter strain were isolated from mouse brains 3-4 weeks post-infection and stained with *Dolichos biflorus* agglutinin (DBA; blue). Cysts expressed both mCherry and GFP, indicating that the truncated *GRA11A* promoter was also active under bradyzoite conditions. **C)** Unsporulated TgVEG oocysts isolated from a cat experimentally infected with TgVEG reporter strain tissue cysts were imaged by phase and fluorescence microscopy. Unsporulated oocysts lacked detectable GFP or mCherry expression but exhibited the expected UV-excited autofluorescence. (top and bottom panels are from two separate oocysts). **D)** TgVEG reporter strain oocysts were sporulated by aeration and shaking for 6-8 days in 2% sulfuric acid and visualized using fluorescence and phase microscopy. Sporulated oocysts showed clear mCherry fluorescence (indicating that the *GRA11A* promoter was activated during sporulation) but did not express any detectable *GRA1*-promoter driven GFP (top and bottom panels are from two separate oocysts). **E)** Sporulated oocysts were used to infect mice intraperitoneally which is known to cause excystation. Peritoneal exudates were placed into tissue culture and parasites that grew from these preparations expressed by mCherry and GFP. All results shown in panels A–E were reproduced in multiple independent experiments (>3). Created in BioRender. AlRubaye, H. (2026) https://biorender.com/9z4xqjb.

To determine if our engineered strain retained full life-cycle competence, five brains from chronically infected CBA/J mice were fed to a cat. Tissue cysts from those brains expressed both GFP and mCherry (Fig. 1B). Feline fecal samples were collected daily between days 3 and 17 post-infection. Unsporulated oocysts of TgVEG:GFP:mCh lacked detectable GFP or mCherry fluorescence (Fig. 1C), whereas sporulated oocysts exhibited mCherry fluorescence but no GFP fluorescence (Fig. 1D). Sporozoites were excysted from oocysts via mouse infection^3^, and parasites were recovered from the peritoneum by lavage and as expected GFP and mCherry fluorescence were detected in TgVEG:GFP:mCh parasites growing as tachyzoites *in vitro* (Fig. 1D). Because mCherry expression was observed in tachyzoites, bradyzoites, and sporozoites, we hypothesize that constitutive expression results from the absence of regulatory elements in the hybrid GRA11A promoter, which has been proposed to be occupied by MORC^20^, along with the physical proximity of this truncated GRA11A promoter to GRA1 sequences in the plasmid driving GFP (Extended Data Fig.1).

### A single cell atlas of *T. gondii* enteric development

We generated a single-cell atlas of *T. gondii* enteric development from 16,966 parasites isolated from infected feline intestines across two independent experiments (Fig. 2A). These intestine-derived parasites displayed transcriptional signatures that were strikingly distinct from *T. gondii* tachyzoites cultured *in vitro* for 8 days post-excystation (Fig. 2B). As expected, the *in vitro* excysted parasites showed high transcript abundance for canonical tachyzoite markers like SAG1 and SAG2A (Fig. 2C; top panels)^21,22^. In contrast, parasites derived from the cat intestine exhibited significantly higher levels of known merozoite markers like GRA80 and PNP (Fig. 2C; bottom panels) compared to tachyzoites^16,19^. Overall, the gene expression profiles of intestine-derived parasites show many unique transcripts that were distinct from those of tachyzoites as well as overlapping transcripts (Fig. 2D), underscoring both the differences between life stages and the remarkable adaptability of *T. gondii*, which enables it to thrive across various host environments. To identify any differences between the cat samples, we downsampled to account for the differences in the cell counts and performed bulk differential gene expression analysis on the resulting data. Very few genes were of significantly different abundance between the two samples (Extended Data Figure 2) and we found the enteric samples from both cats to be highly correlated (Fig. 2E)

**Fig. 2:**
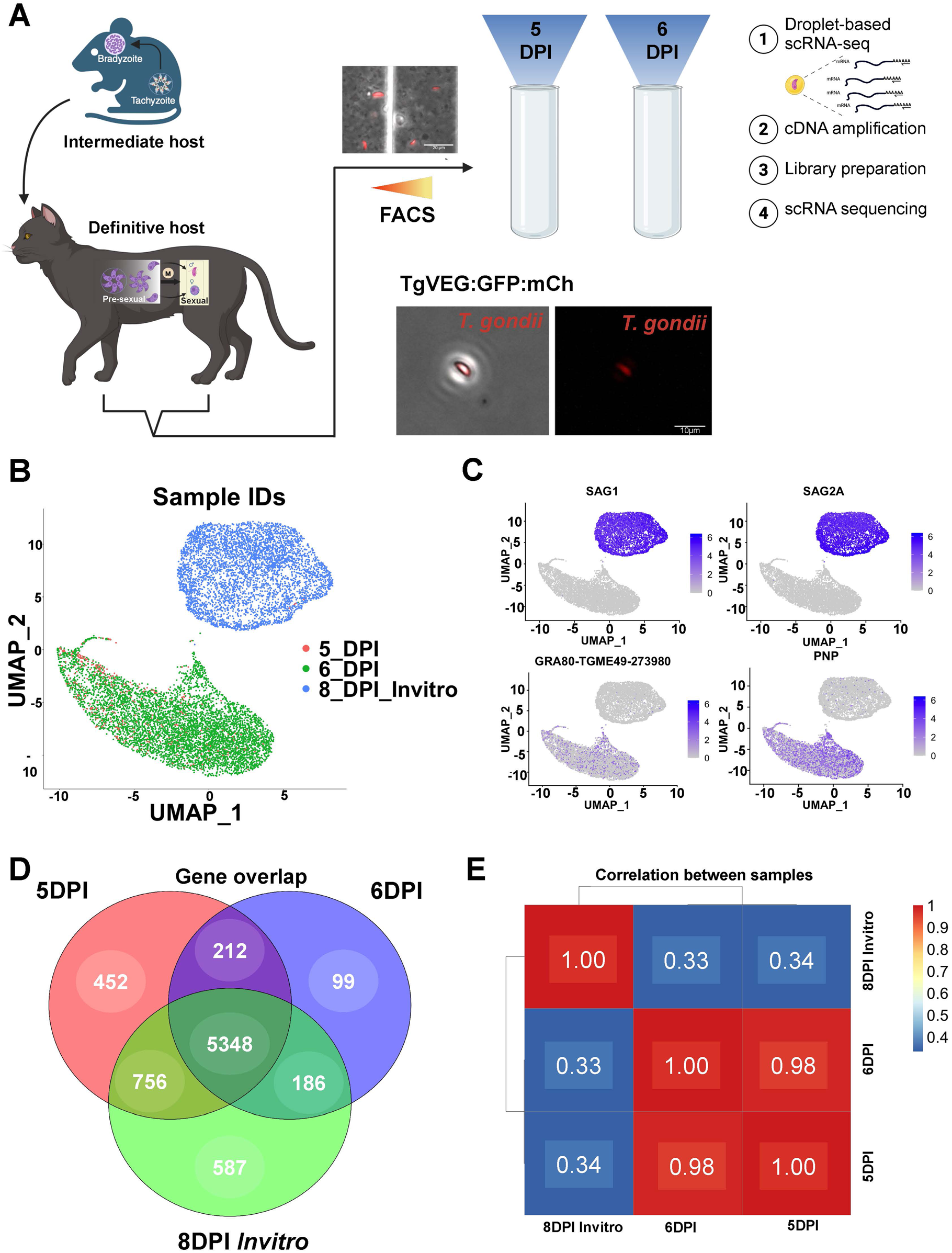
scRNA-seq transcriptome landscape of *Toxoplasma gondii* sexual and asexual development. **A)** Experimental workflow: Mice were infected with the TgVEG reporter strain for 25 days. Brain tissue containing bradyzoite cysts from 28 chronically infected mice was fed to cats (n = 2). Enteroepithelial stages were isolated from the small intestine of a cat euthanized at 5–6 days post-infection (DPI), sorted by mCherry fluorescence, and subjected to single-cell RNA sequencing (10x Genomics). Created in BioRender. AlRubaye, H. (2026) https://biorender.com/bjhm7zo. Images shown are from mCherry+ parasites from a crude, filtered intestinal scraping prep on a hemacytometer (top) and a single mCherry+ parasite detected after sorting (bottom). **B)** UMAP visualization of scRNA-seq data from cat enteric stages (D5 and D6 DPI) and TgVEG tachyzoites cultured for 8 days post-excystation. Distinct clustering reflects the clear and expected transcriptional differences between replicating TgVEG tachyzoites and feline intestinal stages isolated by flow cytometry. D5 had fewer sequenced cells than D6. **C)** Expression profiles of known tachyzoite markers (top two panels; SAG1 and SAG2A) and merozoite markers (bottom two panels; GRA80 and PNP), highlighting stage-specific transcript abundance. **D)** Venn diagram showing overlap of detected transcripts across cells from 5 and 6 DPI taken from the feline intestine, and cells isolated from *in vitro* cultures of TgVEG for 8 days post-excystation from oocysts. **E)** Correlation analysis of transcriptomes across samples, illustrating the degree of similarity between *in vitro* tachyzoites and enteric stages at 5 and 6 DPI and high dissimilarity with *in vitro* cultivated TgVEG.

### Identifying distinct clusters of *T. gondii* in enteric stages

To identify distinct clusters in the feline enteric stages, doublets were detected and removed using scDblFinder algorithm^23^, with the neighborhood parameter pK optimized to 0.08^24^. Artificial doublets were generated from 25% of cells (pN = 0.25), and dimensionality reduction was performed using the top 25 principal components determined by the elbow method (Extended Data Fig. 3B). Predicted doublets were excluded prior to downstream analysis. In addition, cells with fewer than 500 unique molecular identifiers (UMIs) were removed based on the knee point detection in EmptyDrops (Extended Data Fig. 3A,C)^25^. After filtering, 15,068 of 16,966 remained. Gap statistic analysis was then applied to determine the optimal number of clusters in the scRNA-seq dataset^26,27^. This analysis identified K = 10 as the optimal cluster number (Extended Data Figure 3D), which was used to partition cells into transcriptionally distinct subpopulations (Fig. 3A). Feature plots of multiple reported *T. gondii* enteric stage-specific *T. gondii* genes show variation in the distribution of transcript abundance across the UMAP landscape (Fig. 3B).

**Fig. 3:**
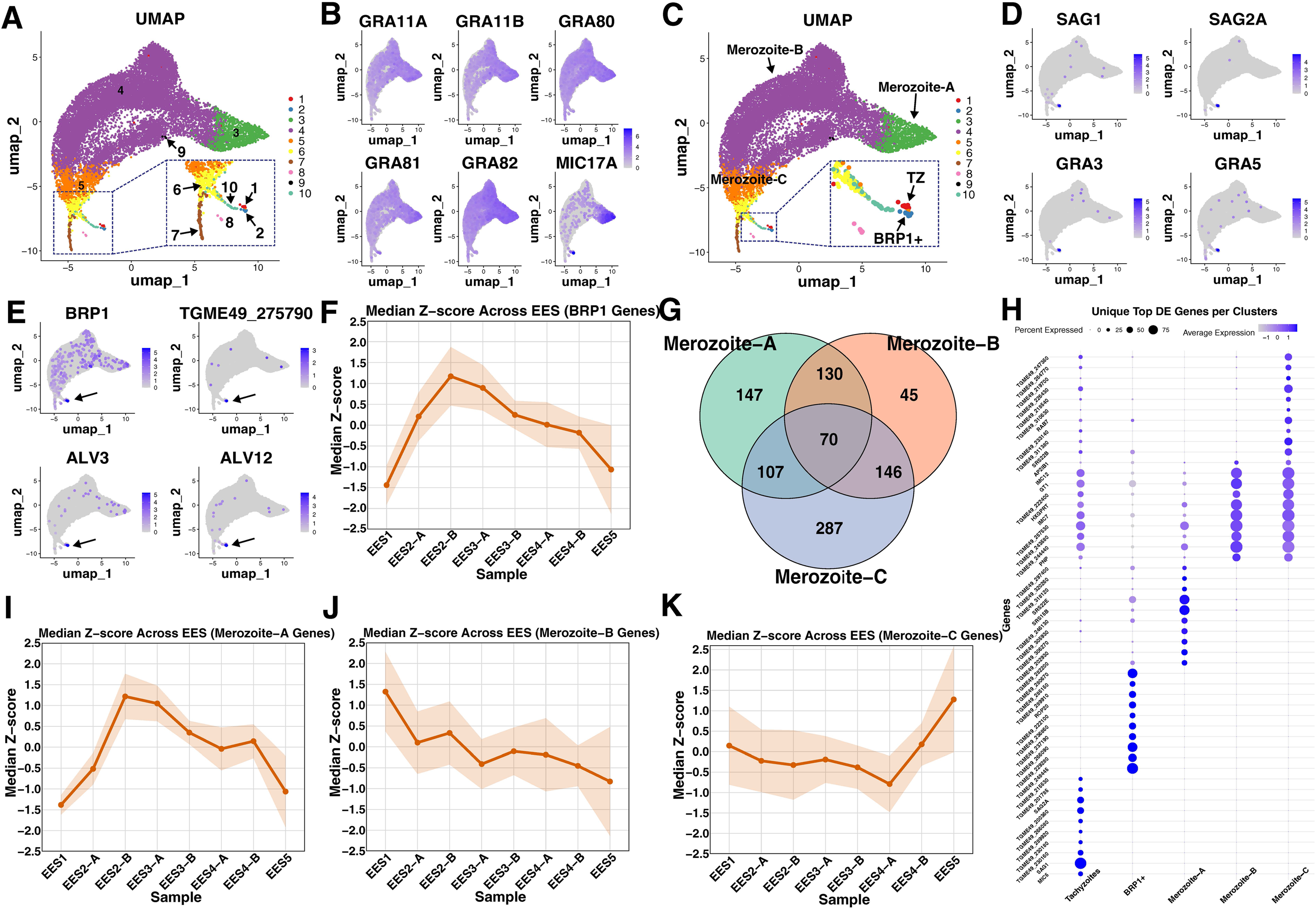
Single-cell transcriptomic profiling of *T. gondii* enteric stages. **A)** UMAP visualization of scRNA-seq data from cat enteroepithelial stages at 5 and 6 days post-infection (n = 15,068 cells). Predicted doublets were removed using scDblFinder. Dimensionality reduction was performed with the top 25 principal components prior to UMAP embedding, and cluster number (10) was estimated using the gap statistic. **B)** Feature plots showing expression of multiple known canonical markers for enteroepithelial stages. **C)** Cluster annotation based on top differentially expressed genes and known markers of merozoites and tachyzoites. **D)** Feature plot of the “tachyzoite” cluster (Cluster 1; arrowhead) highlighting expression of multiple canonical tachyzoite markers. Cat intestine-derived tachyzoites are thought to ultimately exit the intestine and infect cells throughout the body including the brain and muscle. **E)** Expression profiles of genes uniquely expressed in Cluster 2 (annotated as BRP1+; arrowhead). Bradyzoite rhoptry protein 1 (BRP1) is known to be expressed in both bradyzoites and cat enteroepithelial stages. **F)** Line plot of median z-scores for the top 100 genes defining the BRP1+ cluster across enteroepithelial stage samples from Ramakrishnan *et al.* (2019) where bulk RNAseq was used to identify changes in transcriptional profiles during cat enteric development. BRP1 cluster associated genes peak early during cat enteroepithelial development (EES2-3). Shaded ribbons indicate ±1 SD. **G)** Venn diagram of genes expressed within and between the Merozoite-A, -B, and -C clusters. **H)** Dot plot of the top 10 differentially expressed genes for the Tachyzoite (**Tz**), **BRP1+** and Merozoite clusters (**MzA**-**MzC**). Dot size indicates the proportion of cells expressing the gene; color intensity reflects average expression. **I–K)** Line plots of median z-scores for the top 100 genes defining the Merozoite-A, -B, and -C clusters across bulk RNAseq samples from Ramakrishnan *et al.* (2019). Merozoite-A associated genes have peak expression in EES1-2 samples from that dataset, while Merozoite-C peaks in the EES4-5 samples that are thought to be enriched for stages later in enteroepithelial development. Shaded ribbons represent ±1 SD. These data suggest a trajectory of development from Merozoite-A to Merozoite-C moving towards gametogenesis.

To assess if any of the clusters were associated with previously reported canonical marker genes, we examined the differentially expressed genes across all clusters and associated them with cell-type based on previous studies. Cluster 1 showed enriched expression of canonical tachyzoite markers like SAG2A and SAG1 (Fig. 3D). Both genes were found to be significantly enriched and uniquely expressed in this cluster (log_2_FC >15; adj. *p* = 0.0). Therefore, we annotated cells in cluster 1 as ‘enteric tachyzoites’ (Fig. 3C). It is possible that these parasites were in the process of dissemination from the intestinal lining to other tissues within the host, since extraintestinal development of *T. gondii* occurs even in the definitive host. Cluster 2 was defined by high expression of genes like BRP1 (log_2_FC >9; adj. *p* = 1.43e-93; Fig. 3E) a bradyzoite and merozoite marker, and AMA2 which is a protein involved bradyzoite invasion (log_2_FC >9; adj. *p* = 1.43e-93; ^5,28,29^). While BRP1 expression was expected in merozoites^30^, it was somewhat unexpected to be restricted to cluster 2 rather than being more broadly distributed. TGME49_275790 (Fig. 3E) is a merozoite-associated gene that is also enriched in this cluster (log_2_FC >11.5; adj. *p* = 0), along with other tachyzoite-associated genes like *ALV3* and *ALV12* (log_2_FC >12; adj. *p* = 0; Fig. 3E). The presence of bradyzoite/merozoite and tachyzoite markers in these cells distinguishes this cluster from the rest of the enteric parasite population and suggests that these parasites may represent an intermediate state between known canonical stages. Therefore, this cluster was annotated as “BRP1+” based on one of its defining markers (Fig. 3E). We compared the transcript abundance of the top 100 genes in the BRP1+ cluster to bulk RNAseq transcriptome data from *T. gondii* cat infections previously published by Hehl and colleagues ^17^ and found that these genes were most characteristic of EES2 and EES3 stages (Fig. 3F).

Clusters 3, 4, and 5 were enriched for multiple known merozoite-associated transcripts (Fig. 3G) including SRS15B, SRS22E, SRS22D, GRA11A, and GRA11B^16,17,19,20,31^. Cluster 3 comprised 1,317 cells and was defined by high expression of TGME49_287040 (log_2_FC = 5.91; adj. *p* = 0) and TGME49_200270 (log_2_FC = 5.87; adj. *p* = 0), transcripts encoding hypothetical proteins that were previously detected in enteric-stage parasites^17^. Cluster 4 was the largest cluster, containing 12,051 cells (∼80% of all cells), and was characterized by the expression of genes like IMC7 (log_2_FC = 1.04; adj. *p* = 0) and TGME49_244440 (log_2_FC = 1.73; adj. *p* = 0)^17^. Cluster 5 is made up of 1,199 cells with multiple differentially expressed transcripts including TGME49_311720 (log_2_FC = 1.8; adj. *p* = 0) and TGME49_293870 (log_2_FC = 1.8; adj. *p* = 0)^17^. Although all three clusters shared core merozoite markers like GRA11A, GRA11B, and GRA82, each exhibited unique gene signatures that distinguish them from one another. The top 10 differentially expressed genes for each cluster are shown in Fig. 3H. Based on these profiles, clusters 3, 4, and 5 were annotated as “Merozoite-A,” “Merozoite-B” and “Merozoite-C” respectively. These designations are distinct from the morphological stages (Type A–E schizonts), which reflect structural characteristics, whereas our annotations are based on transcriptional profiles.

Cluster 6 was enriched for transcripts associated with late-stage development, as defined by comparative expression analysis with data from Ramakrishnan *et al*^17^. Among the top differentially expressed genes were TGME49_266960 (log_2_FC = 3.38; adj. *p* = 0) and TGME49_290200 (log_2_FC = 4.41; adj. *p* = 0; Fig. 4B). When compared to Ramakrishnan *et al*^17^, the top 100 genes in this cluster exhibited increased expression in late EES samples (EES4 and 5; Fig. 4C). Based on data presented immediately below, we hypothesize that cluster 6 represents a pre-gamete stage of development. Therefore, this cluster was annotated as “Pre-gamete” (Fig. 4A).

**Fig. 4:**
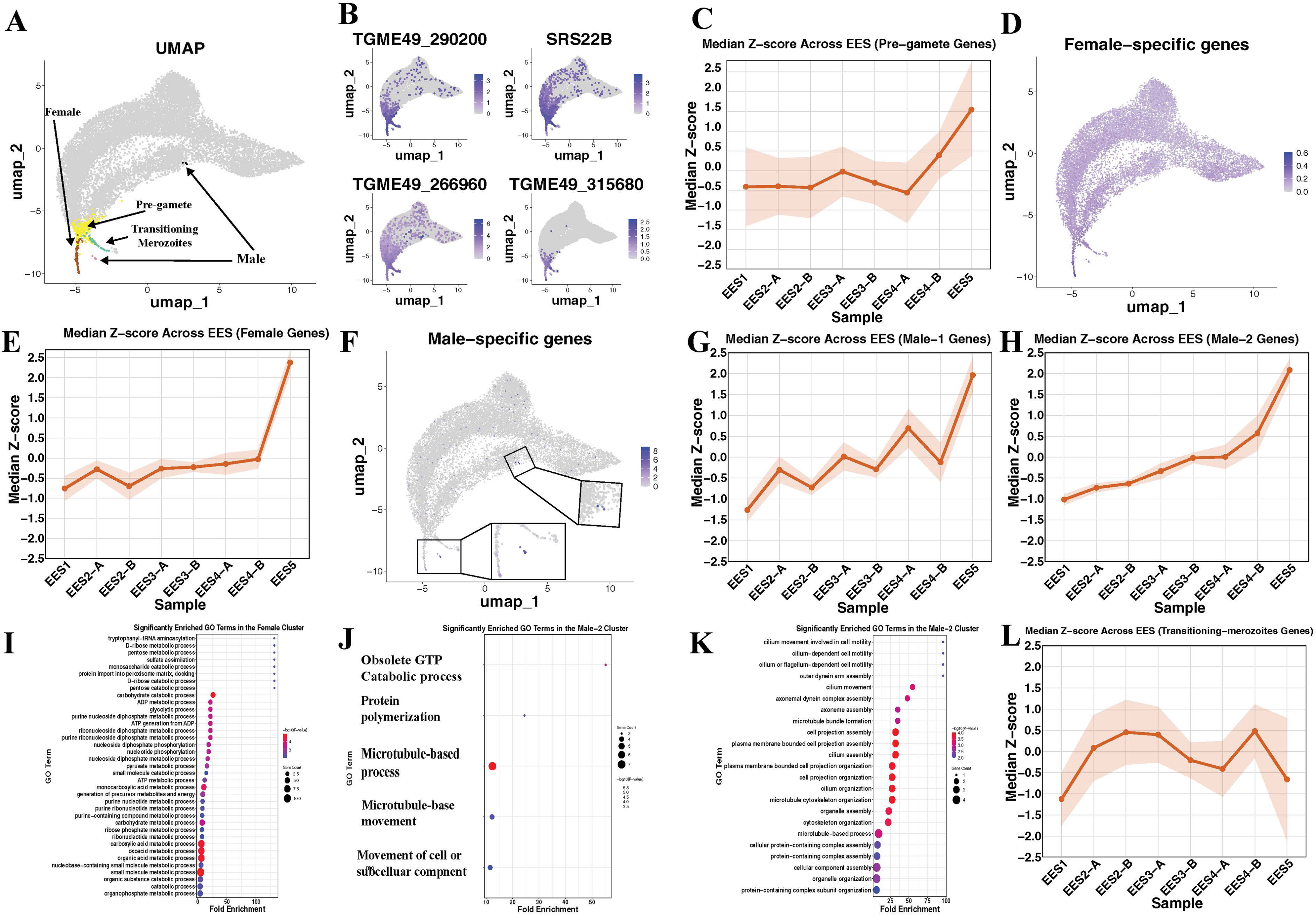
Transcriptomic signatures of pre-gamete and gamete-associated clusters in *T. gondii*. **A)** Annotation of Clusters 6–10 based on top differentially expressed genes. **B)** Feature plots showing genes enriched in the pre-gamete cluster. **C)** Median z-scores of the top 100 pre-gamete cluster genes across EES samples from Ramakrishnan *et al.* (2019). Lines represent median expression per sample; shaded ribbons indicate ±1 SD. **D)** 106 female-specific genes identified in *T. gondii*, *Plasmodium berghei*, and *Cryptosporidium parvum* exhibit transcriptional enrichment in cluster 7. **E)** Median z-scores of the top 100 female cluster genes, showing increased transcript abundance in late EES samples. Shaded ribbons indicate ±1 SD. **F)** Distribution of male-specific genes across clusters. Feature plot of 46 male-specific genes identified by Ramakrishnan *et al.* (2019) shown to be enriched primarily in clusters 8 and 9. **G–H)** Median z-scores of the top 100 genes defining the Male-1 (cluster 8) and Male-2 (cluster 9) clusters, respectively. Both clusters show increased expression in later EES samples from Ramakrishnan *et al.* (2019). Shaded ribbons indicate ±1 SD. **I–K)** Gene Ontology (GO) enrichment analysis of female (cluster 7), Male-1 (cluster 8), and Male-2 (cluster 9) clusters. Female cluster terms are significantly associated with multiple metabolic and biosynthetic processes; both male clusters have genes that enriched for functions related to cellular movement (cilia, microtubule-based movement and organization, axoneme assembly). GO enrichment was calculated in ToxoDB using Fisher’s exact test (one-sided) with Benjamini-Hochberg or Bonferroni corrections for multiple testing. **L)** Median z-scores of the top 100 genes defining the transitioning merozoite cluster in relation to the bulk RNAseq data from Ramakrishnan *et al.* (2019). Shaded ribbons indicate ±1 SD.

To determine if clusters contained female gametocytes, we examined HOWP1 and AO2, known female markers in *T. gondii*, which were exclusively expressed in cluster 7 (Extended Data Fig 4A and 4B)^32^. Furthermore, we used comparative transcriptomics between recently published *Cryptosporidium parvum*, *Plasmodium* and our datasets to identify conserved female-specific markers^33^. Out of 108 female gametocyte-specific genes identified across *T. gondii*, *C. parvum* and *P. berghei*^32–34^, we found that 28 of them were enriched in Cluster 7 (Fig. 4D; Extended Data Fig 4C; Supplementary Table 1), including *TGME49_272900*, whose *Plasmodium* ortholog (*PF3D7_0816800)* encodes the female-specific meiotic recombination protein DMC1^35^, and *TGME49_294820*, (a type I fatty acid synthase) a protein found in high abundance in oocysts from *T. gondii* which is also enriched in *Eimeria tenella* gametocytes^36^. The expression of reported female-specific genes supported the annotation of this cluster as ‘Female’, thereby providing the first detailed transcriptional view of *T. gondii* macrogametocytes. Among the top 20 genes in the female cluster, three belong to the family D protein family (TGME49_236975, TGME49_316670, and TGME49_277572), a large family of tandemly duplicated genes of mostly unknown function. Additionally, we found that a putative surface antigen, SRS46, is uniquely expressed in these cells making SRS46 a candidate surface marker of macrogametocytes. Furthermore, we examined the top 100 differentially expressed genes with the cluster and found that their transcript abundance was increased in late developmental samples from Ramakrishnan *et al*^17^ (Fig. 4E), consistent with the timing of macrogamete development.

Similar to macrogametes, very little is known about gene expression in *T. gondii* microgametes. To identify these cells in our scRNAseq-derived clusters, we used putative microgamete markers from Ramakrishnan *et al.* including genes involved in axoneme assembly, flagellar transport, cell motility, and fertilization^17^. Mapping these genes onto our scRNAseq projection (Fig. 4F; Supplementary Table 2) revealed a small number of cells across two clusters (9 cells in cluster 8 and 3 cells in cluster 9) that express microgamete marker genes. We recognize the potential limitation of identifying so few cells with such characteristics; however the relative rarity of microgametocytes compared to macrogametes is consistent with the literature^37,38^. The enrichment of previously identified male-specific genes (see below) in these clusters supported their annotation as ‘Males-1’ and ‘Males-2’, respectively^17^. Furthermore, we observed a similar female-biased ratio in a subsequent scRNAseq experiment from cat enteroepithelial stages, where we identified 11 male and 106 female cells (Extended Data Fig. 6). The Male-1 cluster expresses Hapless 2 (*HAP2*; TGME49_285940), a male-specific gene essential for gamete fusion and subsequent fertilization in both *T. gondii and Plasmodium* (Extended Data Fig. 5A)^17,39,40^. In contrast, the Male-2 cluster exhibits higher expression of genes related to motility, such as *sperm associated antigen 6* (or *PF16;* TGME49_297820), as well as genes encoding for flagellum-associated proteins (Extended Data Fig. 5B). The *PF16* protein has been observed to associate with the growing cytoplasmic axonemes along the emerging flagella in *T. gondii*^20^, and in *P. berghei*, PbPF16 is expressed in the male gametes^41^. PbPF16 is also essential for maintaining the correct microtubule structure in the axoneme’s central apparatus. This shows that PbPF16 plays a critical role in flagellar formation, a process essential for male motility. Hence, it is possible to speculate that the Male-1 cluster represents intracellular stages (either from a developing macrogamont or a microgamete that has invaded a cell in search of the macrogamete), and that the Male-2 cluster consists of extracellular microgametes, potentially searching for host cells with macrogametes. When we compared the top 100 genes from clusters 8 and 9 to the transcriptomic data reported by Ramakrishnan *et al*^17^., we observed a strong concordance with genes enriched in samples from later developmental stages (Fig. 4G and 4H), consistent with the expected timing of microgametocyte development.

Another branch emerging from the trifurcation point of the pre-gamete cluster contains cluster 10. Marker genes that are significantly enriched within cluster 10 include SRS48Q (log_2_FC > 6.7; adj. *p* = 9.22E-164) and SAG2D (log_2_FC > 8.23; adj. *p* = 0), which are known to be associated with merozoites^17^. Although cluster 10 shares some of the same genes with the Female and Males-1 clusters, such as TGME49_294640 and TGME49_211030, it lacks the expression of sex-specific genes. Specifically, SRS46, AP2X6, and TGME49_293368 are not detected in this cluster. The expression profile of cluster 10 may indicate that these are cells re-entering the asexual cycle (merogony) after failing to differentiate into macrogametes or microgametes. This could suggest a dynamic process where cells have the potential to shift between pre-gamete and merogony depending on specific regulatory signals. Therefore, we have named this cluster “Transitioning-merozoites” reflecting its intermediate state between pre-gametes and merozoites (Fig. 4A). Furthermore, the top 100 genes from cluster 10 do not show an obvious association with EES1-5 from Ramakrishnan *et al.* (Fig. 4L), suggesting that this is potentially a transient population. We did not detect oocysts in our dataset, likely because they lacked fluorescent markers and were therefore not sorted. Additionally, the oocyst wall is highly resistant to chemical disruption, making it plausible that intact oocysts did not lyse during single-cell preparation, which would prevent RNA release and detection.

To infer a developmental trajectory, we performed pseudotime analysis using Monocle3^42^. Because the precise root population initiating enteric development is unknown, we defined the trajectory root by inverting the gamete clusters, positioning the earliest transcriptional states opposite to gametes. This approach placed the Merozoite-A cluster at the beginning of the pseudotime axis (Extended Fig. 7D). The pseudotime trajectory aligned with the expected direction of development, with cells progressing from merogony toward gametogenesis.

We expanded our analysis to investigate the biological processes distinguishing identified male and female clusters by using GO term enrichment analysis on ToxoDB. The top 50 genes in the female cluster were enriched in GO terms such as ‘carbohydrate catabolic process’ (GO:0016052), ‘small molecule metabolic process’ (GO:0044281), and ‘carboxylic acid metabolic process’ (GO:0019752) suggesting unique changes in specific metabolic pathways (Fig. 4I). It is possible that female gametes may rely more on carbohydrate and carboxylic acid catabolism, as well as small-molecule metabolic pathways, potentially to support ATP production and biosynthetic processes required for fertilization and oocyst development. In contrast, the top 50 genes in the Male-1 cluster revealed overrepresentation of GO terms related to ‘microtubule-based movement’ GO:0007018’ ‘microtubule-based process’ (GO:0007017), and ‘protein polymerization’ (GO:0051258) (Fig. 4J). The prominent GO term, ‘microtubule-based process,’ similar enrichment was reported in *P. falciparum*^43^ microgametes, suggesting a potential role for these genes in shaping microgametes or driving the motility of flagellated male gametes. Furthermore, GO term enrichment analysis of the top 50 genes from the Male-2 cluster revealed enrichment in terms such as ‘cilium assembly’ (GO:0060271), ‘plasma membrane-bounded cell projection assembly’ (GO:0120031), and ‘cell projection assembly’ (GO:0030031), emphasizing the important role of structural and motility-related processes in this male cluster (Fig. 4K).

### *In vivo* Perturb-seq screening identifies AP2X6 as a putative regulator of asexual enteric development in the cat

To genetically identify novel regulators of merogony and sexual commitment, we conducted a targeted genetic screen, selecting 28 candidate genes based on their distinct transcriptional profiles observed in our scRNA-seq dataset (Extended Fig. 4A). These genes encode DNA- or RNA-binding proteins, surface antigens, and hypothetical proteins, and, more importantly, 23 out of 28 exhibit unique expression profiles in sexual stages. Three different protospacer sequences per gene were cloned into a *T. gondii* Perturb-seq vector using Gibson assembly and validated by whole plasmid sequencing. The validated plasmids were pooled and transfected into TgVEG:GFP:mCh parasites, generating a knockout pool that was used to infect HFFs (Fig. 5A). To minimize bottleneck effects, each gene knockout was propagated as an independent population, resulting in 28 distinct knockout parasite populations (Extended data Fig. 8A). These populations were individually inoculated into mice at doses of 1,000 or 10,000 parasites per mouse (6–8 mice per population). At 25 DPI, brain tissue from chronically infected mice was divided into equal portions (∼16 g each) and fed to two cats. Parasite shedding was monitored, and upon detection parasites were isolated from the small intestine by scraping and cell sorting for fluorescent parasites at 6 and 7 DPI (Fig. 5A).

**Fig. 5:**
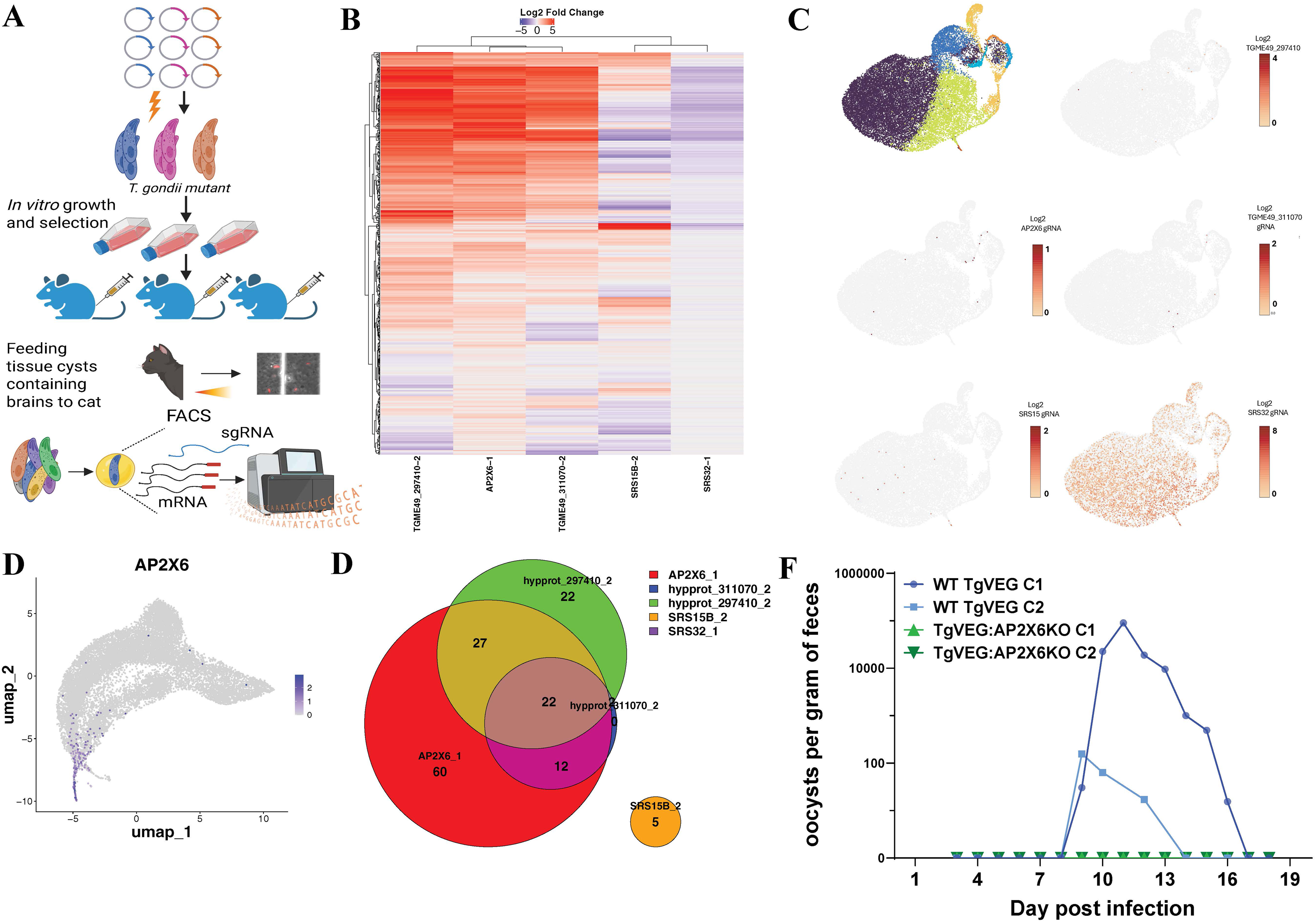
Perturb-CRISPR screen and transcriptomic impact of candidate gene disruptions in *Toxoplasma gondii*. **A)** Schematic of the perturb-CRISPR workflow. For each gene in the 18-candidate list, three distinct gRNAs targeting the same gene were transfected into TgVEG parasites to minimize bottleneck effects. At 24 h post-infection, cultures were switched to cDMEM containing 3 μM pyrimethamine for 12 days after which each knockout population was injected into 6–8 mice. After 25 days, brains were harvested for DNA extraction and for feeding to two cats. At 6–7 days post-feline infection, enteric stages were isolated from cat intestines, syringe-lysed, filtered (40 μm), and sorted by fluorescence followed by perturb-seq analysis. Created in BioRender. AlRubaye, H. (2026) https://biorender.com/3vb2wij. **B)** Heatmap of gRNAs detected within individual cells and their association with significant (|log2 Fold change|>1; Padj<0.01) changes in the abundance of specific transcripts based on analyses within the LOUPE browser (10X Genomics). **C)** UMAP visualization and cluster membership of single cells from the perturb-seq experiment (upper left), followed by panels showing which cells were found to express the 5 gRNAs that were associated with significant changes in transcript abundance. **D)** Feature plot showing transcript abundance of *AP2X6*, a putative *T. gondii* transcription factor that is associated with the female cluster. **E)** Venn diagram illustrating shared and unique transcripts among cells harboring one of the 5 gRNAs that were associated with significant transcriptional changes. Only transcripts with consistent directional changes (all upregulated or all downregulated across compared groups) were included. **F)** Daily oocyst counts from cats infected with WT *T. gondii* VEG (N=2) or *T. gondii* VEGΔAP2X6 (N=2). Cats infected with WT parasites had detectable oocyst shedding as early as D9 PI and shed a total of 5.35 million (C1) and 8000 (C2) oocysts. Cats infected with AP2X6_KO parasites shed no detectable oocysts over the entire course of the experiment. All cats were confirmed to be seronegative for *T. gondii* prior to infection and seropositive for *T. gondii* after infection (see supplementary data).

We assessed library complexity by scRNA-seq of transfected parasites post-selection, detecting protospacers in 23,285 of 37,729 cells (∼62%) with 22.3% sequencing saturation, consistent with a highly complex library even at the low sequencing depth using the 10X genomics system. We assessed transcriptional changes *in vitro* using differential gene expression (DGE) analysis across target gene populations but found no significant differences, consistent with the expression profile of the targeted genes, which are predominantly active during the feline enteric stages and poorly expressed in tachyzoites.

In the cat Perturb-Seq sample, 5,060 cells harboring single gRNAs targeting 18 different genes were recovered (∼19.5% of cells). This is lower than the expected recovery of cells harboring gRNAs, which could be influenced by multiple factors including a bottleneck within the cat small intestine. Cells expressing multiple gRNAs were filtered out, leaving 4,741 cells with single gRNA sequences for downstream analysis. We found 5 specific perturbations in individual cells that resulted in significant transcriptional changes in multiple genes (Fig. 5B). AP2X6 perturbation led to significant increases in transcript abundance for 121 genes relative to parasites with other gRNAs, causing the largest perturbation of the parasite transcriptome across all detected gRNAs. In cells harboring the SRS32-1 gRNA, we observed a significant decrease in transcript abundance in 79 genes (Fig. 5B). Cells carrying the TGME49_311070-2 gRNA, a homolog of *fd1* in *Plasmodium falciparum* 3D7^34^ and associated with a female gametocytes (Extended Data Fig. 7A), had significant transcriptional changes in 36 genes. Furthermore, cells captured with TGME49_297410 and SRS15B-2 gRNAs were found to have changes in transcript abundance in 73 and 5 genes, respectively. Cells harboring the gRNA corresponding to significant perturbations are mapped on the UMAP (Fig. 5C). Some perturbations were observed to impact the same group of genes, exposing a potential link in their function (Fig. 5E). However, follow-up experiments are required to investigate the functions of these genes during enteric development and to confirm that they participate in similar or connected gene regulatory networks.

To assess the role of AP2X6 in sexual development, we generated two TgVEGΔAP2X6 clones and a passage-matched control, confirmed by whole-genome sequencing. Eight mice per strain were infected, and tissue cysts collected at 22 DPI were fed to cats (n = 2 per group). Cats infected with TgVEG shed oocysts at 9 DPI, whereas those infected with TgVEGΔAP2X6 did not (Fig. 5F; Table 5). All cats were seronegative at 0 DPI and seropositive at 24 DPI, as validated by IFA (Table 5). These findings demonstrate that AP2X6 is required for oocyst production and/or release, potentially by repressing merozoite gene expression to permit female gamete development.

## Discussion

Enteric stages of *T. gondii* are difficult to study due to their restriction to feline hosts and challenges in isolating parasites from intestinal epithelial cells. Traditional bulk RNA sequencing approaches have been used on cat enteric stages^16,17^ but these bulk studies do not capture the heterogeneity and asynchronous development. In recent years, single-cell transcriptomics has emerged as a powerful tool, addressing these challenges and enabling characterization of complex developmental trajectories in a wide array of organisms including apicomplexan parasites^33,43–45^. In this study, we developed a constitutively active reporter system using a hybrid promoter, enabling FACS-based isolation of enteric-stage parasites from feline intestine. We then performed scRNA-seq and computational analysis to define the transcriptional program underlying feline-specific development. Using canonical markers, we identified 10 transcriptionally distinct clusters, including tachyzoites, merozoites, transitional forms (pre-gametes), and gametocytes. Gametocyte clusters were enriched for late-stage markers (i.e., EES5 samples in Ramakrishnan *et al.*) and supporting the late developmental identity of these clusters^17^.

One of the most striking observations from our dataset is the disproportionate representation of parasites undergoing merogony compared to those committed to sexual development. Gametocytes were significantly outnumbered by merozoites, suggesting that only a small fraction of parasites proceed toward gametogenesis. However, we acknowledge that this estimate may be influenced by technical and biological factors. Specifically, FACS gating may have excluded pre-gamete or gamete cells with low or absent mCherry expression, and the sampled time points may not have captured peak gamete abundance. Our data support a model in which some pre-gametes may re-enter the merogony cycle, forming what we term “transitioning merozoites.” Our observations suggest that not all cells within the cat enteric environment progress to gametogenesis. Based on clustering and marker expression profiles, we hypothesize that the majority of cells remain in an asexual enteric replication state rather than committing to sexual development. This indicates that gamete formation may be a restricted or regulated process, potentially requiring specific cues or conditions that are not detected by the majority of *T. gondii* cells during development in cat enterocytes. Such plasticity highlights the dominance of asexual replication in apicomplexan life cycles and raises important questions about the regulatory mechanisms that govern stage transitions. This skewed distribution is consistent with observations in *P. falciparum*, where in controlled human malaria infections, fewer than 10% of parasites commit to gametocyte formation^46,47^. It is reasonable to speculate that by maintaining a large comparatively large asexual population it can sustain infection and prolong the window of opportunity for mating, maximizing transmission potential. This strategy may be evolutionarily conserved across apicomplexans.

Although our dataset constitutes significant progress towards gaining an understanding of the gene expression dynamics in *T. gondii* enteric development, the comparatively low number of male gametes indicate that we have yet to fully determine the spectrum of transcriptional profiles in this life stage. This low number compared to female gametes is consistent with expectations based on observations in *P. falciparum*^48,49^. Some male gametes may also be motile and either invade host cells rapidly or be swept away in the cat gut and/or when we wash the intestine prior to scraping. However, there is little doubt that these small clusters of cells represent male gametes. For example, the centrin family protein (TGME49_237490) is associated with the basal structure and function of flagella^50^, sperm-associated antigen 6 (TGME49_297820) is a conserved marker of male gametes, and *HAP2* is a well-characterized protein required for gamete fusion during fertilization^51,41,20,17,32^. The specific and exclusive expression of these genes in the annotated male gamete clusters supports their annotation, despite their comparative low numbers. To address the challenge of low cell abundance, future studies could use male-specific markers identified in this study to enrich for male gametocytes during isolation, allowing for deeper sequencing and potentially functional studies with these poorly understood life stages.

Through scRNA-seq, we identified 28 candidate genes and used Perturb-seq to assess their roles in enteric development. Among the genes with significant transcriptional impact was TGME49_311070, which is a female-specific gene. TGME49_311070 encodes a putative RNA-binding protein and is a homolog of the fd1 gene in *P. berghei*, which, when knocked out, results in infertile females^34^. AP2X6-targeted cells showed higher abundance in transcripts for genes expressed earlier in development (prior to female gamete formation). Therefore AP2X6 could be a transcriptional repressor of merozoite genes, a role consistent with most functionally characterized AP2 factors in *T. gondii*^19,52–54^. AP2X6-targeted cells had higher abundance of genes associated mostly with the Merozoite-A and BRP1+ clusters. However, given the low number of detected cells harboring AP2X6 gRNAs in our perturb-seq experiment, it is possible that the differences in transcript abundance are because all AP2X6-gRNA harboring cells are found outside of the female gamete cluster. Regardless, AP2X6 is clearly required for oocyst production in *T. gondii*, consistent with a potential role in female gamete formation. Cats infected with TgVEGΔAP2X6 failed to shed oocysts despite being seropositive, indicating that loss of AP2X6 disrupts gametogenesis and oocyst formation. Collectively, these findings position AP2X6 as being important for oocyst production and given its expression profile it may be a critical regulator of macrogamete differentiation. Future studies should investigate whether AP2X6 is sufficient to drive macrogametocyte formation *in vitro*, particularly in the context of AP2XII-1 and AP2XI-2 knockdown parasites^19^, and use scRNAseq to identify and profile any female gametes in cats infected with AP2X6 KO parasites to determine if they cannot form at all or if they are transcriptionally distinct from wild-type parasites.

In summary, our single-cell data provide valuable insights into the gene expression dynamics during the enteric development of *T. gondii* in cats. This work holds the potential to uncover previously unknown genetic drivers of differentiation, particularly those involved in gametogenesis. These findings contribute to ongoing efforts in the field to induce gametogenesis, mating, and oocyst formation *in vitro*, paving the way for future progress in manipulating the parasite’s life cycle.

## Materials and methods

### Parasite strain and growth conditions

*TgVEG* parasites were used in this study. These were early passage parasites (P6) that were excysted from sporulated oocysts as described^55^, and maintained in human foreskin fibroblast (ATCC CRL-4001 HFF) cultures incubated at 37°C and 5% CO in Dulbecco’s Modified Eagle Medium (DMEM; Gibco) supplemented with 10% fetal bovine serum (FBS) (Atlas Biologicals), 2 mM L-glutamine (Thermo, Fisher Scientific, Waltham, MA), and 50 mg/mL penicillin-streptomycin and cryopreserved in 10% DMSO.

### Plasmid construction

The TgVEG:GFP:mCh reporter strain was generated by introducing a cassette containing a truncated pGRA11A promoter upstream of mCherry into the pClick-GFP-Luc vector^56^. All primers used in this study are listed in Supplementary Table 3. The pTrunGRA11A was amplified from *TgVEG* genomic DNA, and mCherry was amplified from pCas9.T2A.HXGPRT^57^. The amplified products of pTrunGRA11A, mCherry, and linearized pClick-GFP-Luc were combined in a Gibson assembly reaction master mix (NEB E2621S) at a molar ratio of 0.014 pmol. The Gibson reaction was incubated at 50°C for 1 hour. Plasmid sequences were verified by Oxford Nanopore sequencing.

### Generation of a *T. gondii* VEG reporter strain

*TgVEG* was transfected with the reporter cassette using standard approaches. Briefly, parasites were mechanically released by needle passage (25 and 27-gauge needles) and then pelleted (10 minutes at 800 x g). Approximately 2 x 10^7^ parasites were re-suspended in 100µl Amaxa Basic Parasite Nucleofector 1 (Lonza) along with 3µg of circular pSAG1::CAS9-U6::sgUPRT (Addgene plasmid #54467)^58^ and 5µg of the pTrunGRA11A-mCherry-pGRA1-GFP cassette flanked by 20 bp of homology regions to the UPRT locus were co-transfected by electroporation (Supplementary Table 3). Transgenic and FUDR-resistant parasites were obtained by selection with 10 µM FUDR (Sigma-Aldrich, St. Louis, MO). We performed a diagnostic PCR to verify homologous integration and gene deletion.

### Cat infections, oocyst isolation, and excystations

For cat TgVEG:GFP:mCh parasites were released from HHF culture mechanically using 25 and 27-gauge needles. The released parasites were then pelleted at 800 x g for 10 minutes, then resuspended in PBS. Ten female mice (CBA/J from Jackson lab) were infected with 1,000 parasites in 200 ul PBS intraperitoneally (IP). The levels of IFN-γ were used as an indication of parasite infection and burden early in infection. Blood serum from infected mice was obtained via submandibular bleed, centrifuged at 100 × g for 5 minutes, and then diluted at a 1:20 ratio. IFN-γ concentrations were determined using BD OptEIA Mouse IFN-γ (AN-18) ELISA Set (Cat. # 551866) according to the manufacturer’s instructions.

Mice were treated with Sulfadiazine (0.4g/L *ad libitum* in drinking water) from 6-10 DPI to increase survival during the acute phage of infection. Throughout the course of infection, the mice were monitored and weighed regularly. At 21DPI, a mouse was euthanized to determine the cyst burden. The brain was harvested and homogenized in 1 ml of PBS (Sigma) and stained with Biotin-conjugated Dolichos biflorus agglutinin (DBA) (1:500; Vector Laboratories, cat. no. RL-1032) for 1 hour at room temperature. Subsequently, the sample was incubated with streptavidin conjugated to Alexa Fluor 350 or 647. Cysts were visualized and counted using a 40x objective on an Olympus IX83 microscope using cellSens software.

The remaining mice were euthanized, and the deskinned skulls and brains were fed to cats. Feces were collected between days 3 and 17 post-infection and examined for oocyst production using sucrose fecal floats. The collected feces were submerged in water and mixed with a wooden tongue depressor to break down the materials. The emulsified mixture was then filtered through 840 µm and 250 µm sieves to further break down the particles. The filtered liquid was poured into 225 mL tubes and centrifuged at 2,000 × g for 10 minutes. The resulting pellet was resuspended in a 1.56 M sucrose solution (30 mL sucrose per tube per 10 g of feces). The sucrose mixture was centrifuged at 1,180 × g for 10 minutes, and the top 10 mL of the supernatant was collected for oocyst quantification using hemocytometer. This oocyst collection step was repeated at least twice to minimize oocyst loss during processing. The isolated sucrose pellet containing oocysts was mixed with water at five times the volume and centrifuged at 1,180 × g for 10 minutes. The pellet was then resuspended in 2% H_2_SO_4_ and kept at room temperature with shaking for 7 days to allow for sporulation. These samples were stored at 4°C.

To excyst *T. gondii* oocysts, sporulated oocysts were treated with 1N NaOH (at a ratio of 3:5 oocyst volume) to neutralize the 2% H_2_SO_4_ solution, and then washed multiple times with Hanks’ balanced salt solution (HBSS; Life Technologies, 14175145) until the pH was ∼7.2. pH of the solution was measured using pH indicator strips. The oocysts were then pelleted at 1,000 x g for 8 minutes at 24°C and resuspended in PBS to prepare for excystation in mice. CBA/J mice were infected with 100 oocysts orally or IP. On days 4-7 post-infection, mice were euthanized, and peritoneal lavage fluids were collected in PBS. Lavage fluid was palleted by centrifugation, resuspended in cDMEM, and then placed on confluent HHFs monolayers for 18 hours. Media was changed as necessary, and parasites were visualized using light and fluorescence microscopy.

### Single-cell transcriptomics from *T. gondii* cultured *in vitro*

For this experiment, we used parasites that were cultured *in vitro* for 8 days post-excystation. The sporozoites were excysted from oocysts as previously described ^59,60^. Post-excystation, the sporozoites were resuspended in culture media and used to infect confluent monolayers of HHFs, grown at 37°C and 5% CO_2_. The parasites were passaged as necessary. At 8 days post-excystation, the parasites were counted and processed for library preparation using 10x Genomics Chromium technology. The libraries were then sequenced on a HiSeq 4000 platform (Illumina) with a targeted depth of approximately 30,000 reads per cell for the 3’ mRNA library.

### Single cell transcriptomics of *T. gondii* tachyzoites and cat enteric stages

For single-cell transcriptomics, cats were euthanized at 5 and 6-DPI. The small intestine was isolated from the euthanized cats and divided into three segments. The small intestine was then washed with PBS and opened. The parasites were isolated from the small intestine of the cat by scraping the intestinal lining with glass microscope slide (Fisherbrand: cat#12-550-112). The scraped materials were syringe-passaged through 18, 22, 25, and 27-gauge needles, followed by sequential filtration through cell strainers (100 μm and 40 μm, Corning). The strained materials were pelleted at 1,000 x g for 8 minutes and resuspended in PBS with 1% FBS. The *Tg*VEG:GFP:mCh parasites in the scraped materials were then sorted based on mCherry-derived fluorescence expression. Sorting was performed using BD FACSAria. The sorted parasites cells were concentrated by centrifugation at 1,000 x g for 8 minutes at 4°C and resuspended in PBS containing 1% FBS (Sigma). The cells were then processed for library preparation using 10x Genomics Chromium technology. The libraries were then sequenced on a HiSeq 4000 platform (Illumina) with a targeted depth of approximately 50,000 reads per cell for the 3’ mRNA library.

### scRNA-seq data processing

Sequencing reads from the 3’ mRNA libraries were processed using Cell Ranger v8.0.1 (10x Genomics). Reads were aligned to a custom *Toxoplasma gondii* reference genome (ME49, based on release 56), including intronic reads, while retaining all other settings at their default values. The custom library also included mCherry fused to the 3’ end of DHFR and GFP fused to the 3’ end of GRA2, as these were components of the reporter cassette. We sequenced 16,695 cells with 42,871 mean reads per cell; with a median UMI of 561 counts per cell. Cell calling was performed using the two-stage algorithm implemented in Cell Ranger, which distinguishes cell-associated barcodes from background noise such as ambient RNA from lysed cells. The barcode rank plot was used to visualize this separation, and the knee point guided the selection of a UMI threshold to exclude low-quality cells and empty droplets. EmptyDrops was additionally applied to refine cell calling. The resulting UMI count matrices were imported into Seurat v5.1.0 or Loupe Browser v8.0.0 for downstream analysis. A UMI threshold of 500 UMIs per cell was applied to filter low-quality cells. Doublets were identified and removed using the scDblFinder algorithm (V1.20.2). The neighborhood size parameter (pK) was empirically optimized to maximize classification accuracy, and artificial doublets were simulated from a subset of cells. Dimensionality reduction was performed using principal component analysis, with the number of components selected by the elbow method. Clustering analysis was performed using the gap statistics method to determine the optimal number of clusters in R.

### Generation of perturb-seq vector

Capture sequence 1 (GCTTTAAGGCCGGTCCTAGCAAG) was inserted into the pCAS9-T2A-DHFR vector using Q5 mutagenesis (NEB) to generate pCAS9-T2A-DHFR-CS1 (TgPerturb-seq vector) and the perturb-seq vector was verified by nanopore sequencing. To generate the pool of perturb-seq vectors used in the screen, single-stranded DNA (ssDNA) oligonucleotides encoding the protospacer sequences for each gene of interest were generated using EuPaGDT^61^. Three ssDNA were synthesized for each gene with 20 bases of homology to either side of the perturb-seq vector integration site (Supplementary Table 4). Each ssDNA was integrated into the linearized TgPerturb-seq vector (NcoI and PacI) by Gibson assembly (vector: insert = 1:5). The Gibson reaction was incubated for 30 mins at 50 °C.

### Perturb-seq experiment

*Tg*VEG:GFP:mCh parasites were transfected with NotI-HF-digested (NEB) perturb-seq vectors by electroporation using Amaxa Basic Parasite Nucleofector 1 (Lonza). For each transfection, approximately 10^7^ parasites were electroporated with 15 μg of pooled Perturb-seq vectors, composed of three vectors expressing different gRNA targeting the same gene (5 μg each). Twenty-four hours post-transfection, 3 μM pyrimethamine (Sigma-Aldrich, St. Louis, MO) was added to the media to select for integration of the perturb-seq vectors into the genome for 12 days. The diversity of perturb-seq vectors was validated by amplifying the gRNAs, followed by sequencing the amplicons.

Female CBA/J or CD-1 mice were infected intraperitoneally with individual knockout parasite populations (CBA/J, 1,000 parasites; CD-1, 10,000 parasites in 200 μl PBS; 6–8 mice per population)The infected mice were treated with Sulfadiazine (0.2g/L) from 4DPI to 10DPI. The treatment was employed to reduce the parasite burden. They were monitored and weighed regularly throughout the experiments. To assess the cyst burden in the brains of infected mice, two mice were euthanized at 21DPI for analysis. The brains were homogenized in 1 ml PBS and subsequently stained with Biotin-conjugated *Dolichos biflorus* agglutinin (DBA) (1:500; Vector Laboratories, cat. no. RL-1032) for 1 hour at room temperature, followed by incubation with Biotin-conjugated to Alexa Fluor 350 or 647. Cysts were visualized and counted using a 40x objective on an Olympus IX83 microscope using cellSens software.

Upon cyst verification, brain tissues from the remaining mice were collected and fed to two pathogen-free cats. Feces were collected from 3 DPI until oocysts were detected at 5 DPI. On 6 and 7 DPI, the cats were euthanized, and parasites were isolated from the small intestine and sorted as previously described above.

The pooled sorted cells were concentrated by centrifugation at 1,000 x g for 8 minutes at 4°C and resuspended in PBS containing 1% FBS (Sigma). The cells were then processed for library preparation using 10x Genomics Chromium technology. The libraries were then sequenced on a HiSeq 4000 platform (Illumina) with a targeted depth of approximately 45,000 reads per cell for the 3’ mRNA library and 10,000 reads per cell for the CRISPR sgRNA library.

### Perturb-seq data processing

The sequencing reads from the 3’ mRNA and CRISPR sgRNA libraries were processed using Cell Ranger v7.2.0 (10x Genomics). The reads were aligned to both a custom *T. gondii* transcriptome reference and another reference containing protospacer sequences. Intronic reads were included in the alignment, while all other parameters were left at their default settings. Cell Ranger assigned the reads to specific cell barcodes, mapped them to the corresponding genes or protospacers, and counted the unique molecular identifiers (UMIs) for each gene or protospacer within each cell barcode. This generated UMI × cell barcode count matrices, which were downloaded for downstream analysis to identify gRNA-specific perturbations using in the single-cell transcriptome Loupe Browser v8.0.0 and Seurat v5.1.0. UMAP embeddings were generated with the following parameters: number of neighbors = 15, minimum distance = 0.1, and 10 principal components, default settings. In total, we sequenced 26,000 cells with a median depth of 1,996 UMI. 5060 cells were detected with one or more protospacer. For analysis, we filtered cells with multiple protospacers to retains cells with a single protospacer detected, 4741 cells, for further analysis.

## Animal statement

Animal experiments were conducted in accordance with guidelines of the American Veterinary Medical Association. Four to eight-week-old female CBA/J mice (The Jackson Laboratory) and CD-1 (Charles River Laboratories) were housed at the University of Pittsburgh for 3–6 weeks prior to experimentation and maintained on a 12-h light/dark cycle with controlled ambient temperature and humidity. Ten to twenty-week-old cats were purchased from Marshall BioResources; sex was determined by availability. Cats were housed at the University of Pittsburgh for 2–10 weeks prior to use in experiments. Mice were euthanized by controlled CO exposure, and cats were euthanized using Beuthanasia. All animal procedures were approved by the University of Pittsburgh Institutional Animal Care and Use Committee (IACUC), protocol no. 22061169.

## Data Availability

All the relevant data to support the findings of this study are included. The single-cell RNA sequencing (scRNA-seq) demultiplexed FASTQ files have been deposited in the NCBI Sequence Read Archive (SRA) under accession number PRJNA1358975. Processed data matrices and metadata used for downstream analyses are available at Zenodo (https://doi.org/10.5281/zenodo.18372462). Source data are provided

## Code Availability

The code for the analysis is available here: https://github.com/ctsdr/Exploring-Toxoplasma-sexual-development

## Acknowledgement

This work was supported by National Institutes of Health grants T32GM133353 to HSA and R01Al116855 to JPB.

## Contributions

Conceptualization: HAS and JPB.; Methodology: HSA, JPB, SMR, RDS, and NW; Investigation: HSA, JPB, SMR, RDS, and NW; Writing original draft: HAS and JPB; Writing, review & editing: HSA; JPB; Funding acquisition: JPB; Supervision: JPB

