## Extended Data Figures for "A single-cell atlas of *Toxoplasma* sexual development in the feline intestinal tract"

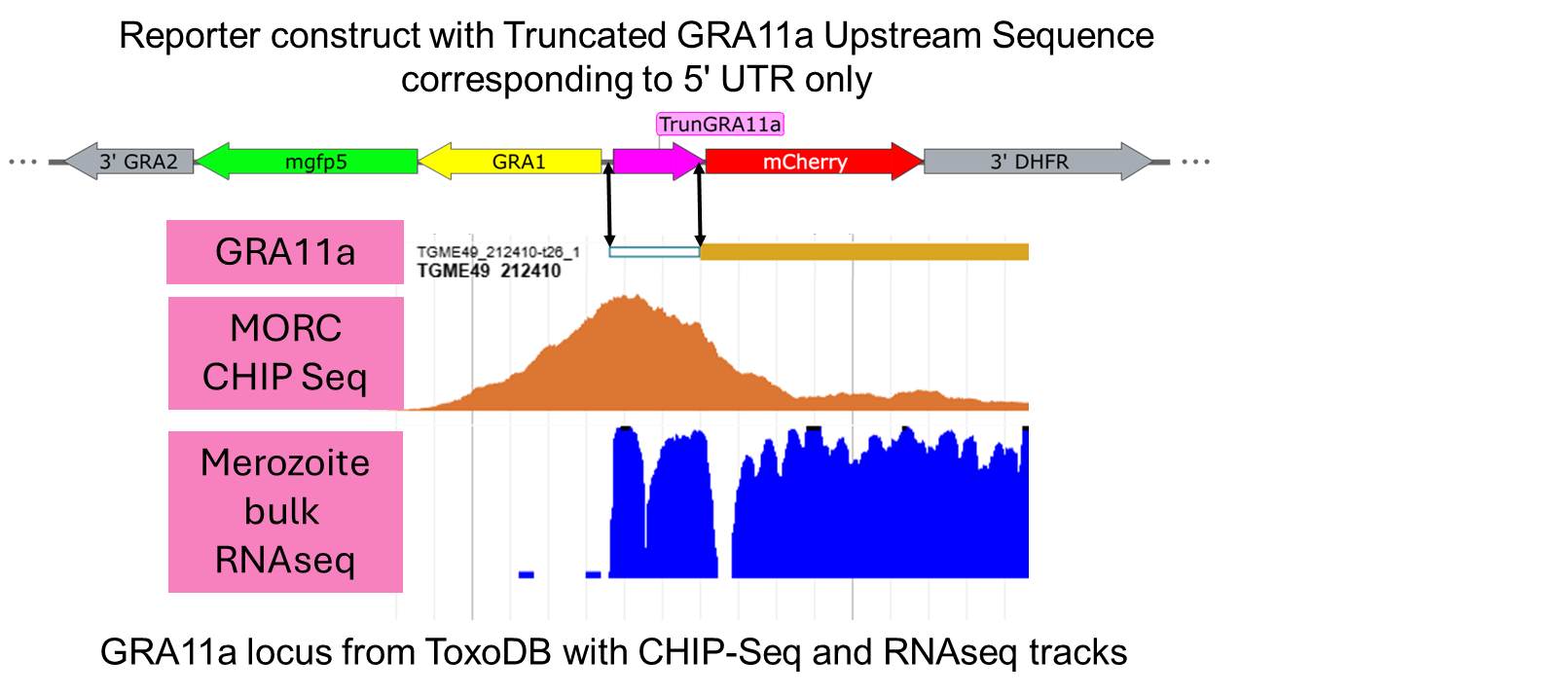

**Extended Data Figure 1:** Schematic of the plasmid containing the reporter gene construct showing all relevant components to scale including the upstream sequence driving mCherry above a scaled alignment between our reporter construct and the GRA11A locus taken from ToxoDB. The GRA11a gene is indicated, and double arrowheads indicate the 5’ UTR of GRA11a as well as the region used in our reporter construct to drive mCherry. MORC and RNAseq tracks are shown which are derived from data published in Farhat *et al.*, Nature Microbiology 2020 and Hehl *et al.*, BMC Genomics 2015, respectively. Both of these datasets are hosted on ToxoDB. The MORC track is named "ChIP-Seq in MORC WT and MORC KD parasites - Pruku80_MORC_HA_Rep1_anti- HA (unique) Coverage" and the RNAseq track is called "Tachyzoite and merozoite transcriptomes - 002.1 - Merozoite (unique forward) Coverage".

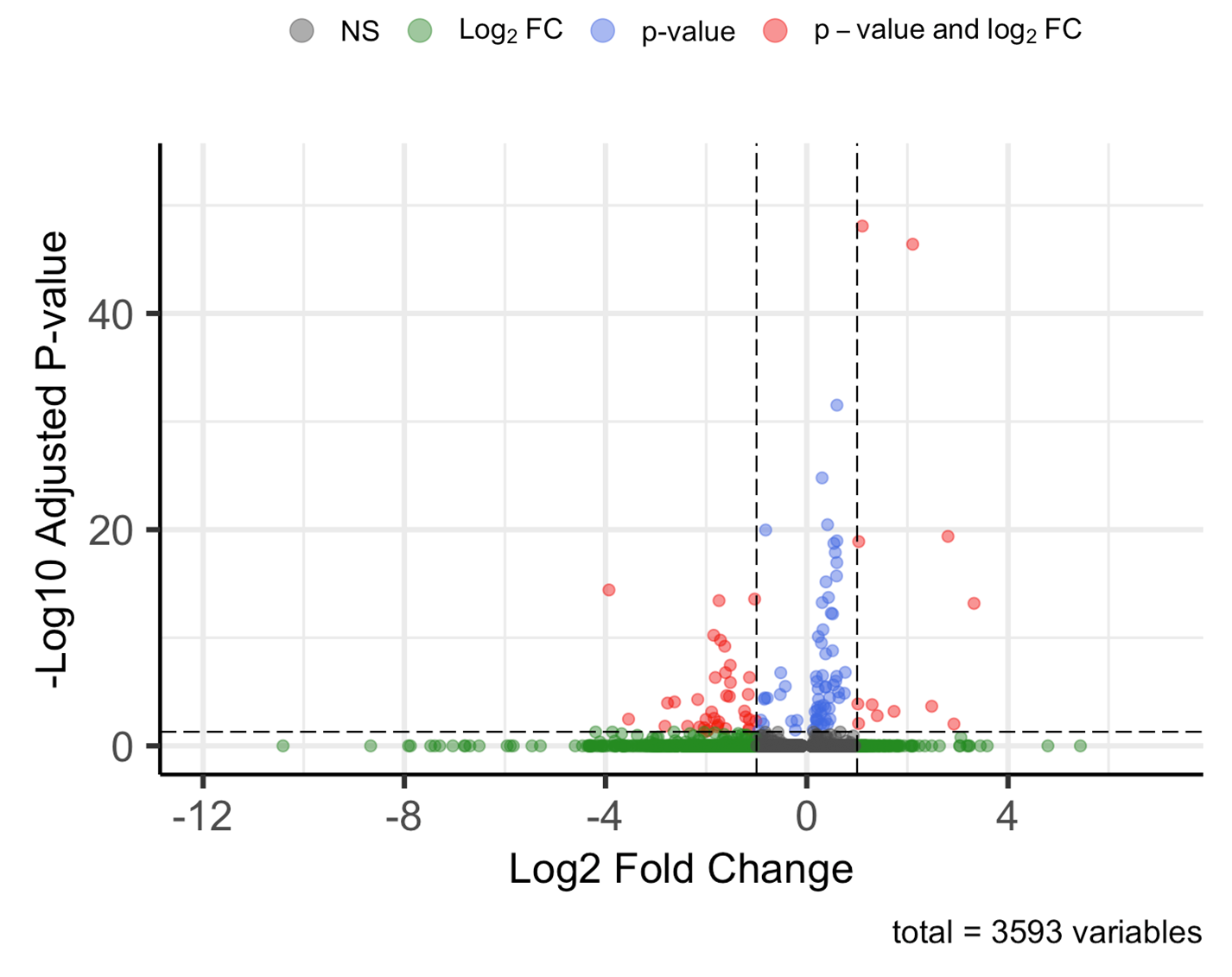

**Extended Data Figure 2:** Enhanced volcano plot illustrating differences in transcript abundance between merozoites isolated at 5 DPI (higher abundance to the left) and 6 DPI (higher abundance to the right). Significantly upregulated and downregulated genes are highlighted in red. The majority of transcripts are of similar abundance in both datasets.

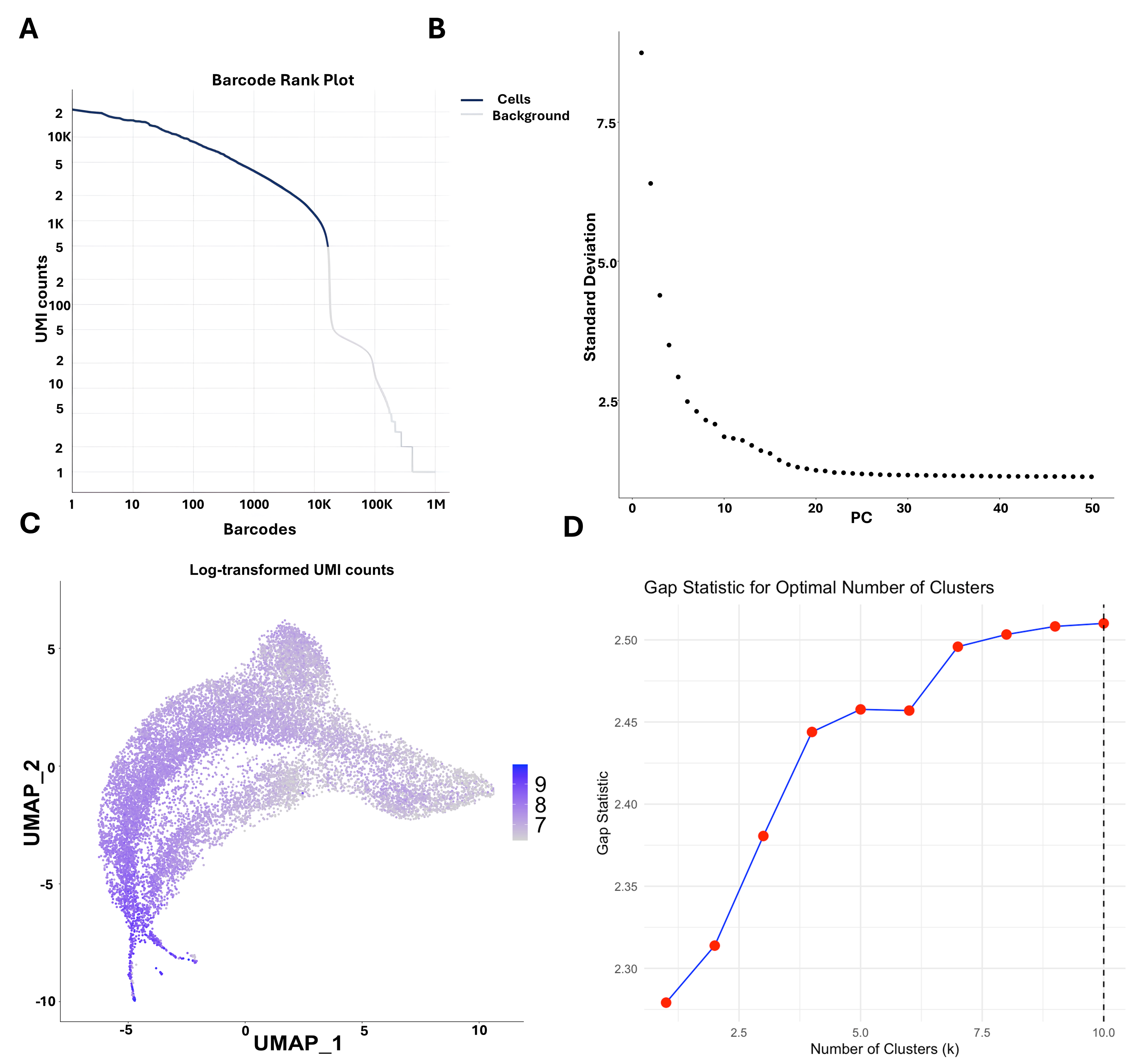

**Extended Data Figure 3:** **A)** Barcode rank plot showing cell calling based on RNA content. The two-stage algorithm distinguishes cell-associated barcodes from background noise, including ambient RNA from lysed or dead cells. **B)** Elbow plot illustrating the total amount of variance attributed to each principal component (PC). **C)** UMIs per cell across the UMAP visualization (Log2-transformed). **D)** Gap statistic analysis used to determine the optimal number of clusters by comparing observed clustering to a random reference dataset. The optimal number of clusters was 10.

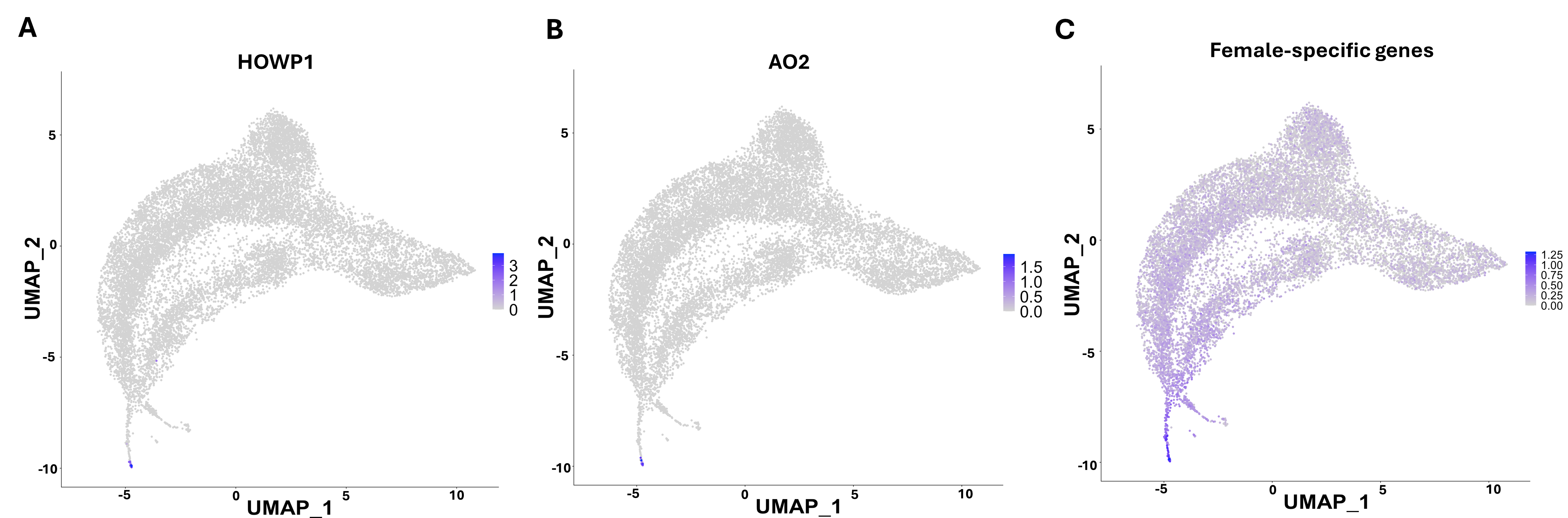

**Extended Data Figure 4:** **A)** Feature plot showing transcript abundance for the *AO2* gene **(A)** or the *HOWP1* gene, two previously identified female-specific transcripts in *T. gondii*. Transcripts for both are uniquely detected only at the tip of where the female cells cluster. **C)** Sum of transcript abundance across 28 of 108 known female-specific genes identified in *T. gondii*, *Cryptosporidium parvum*, and *Plasmodium berghei* which are enriched in cluster 7.

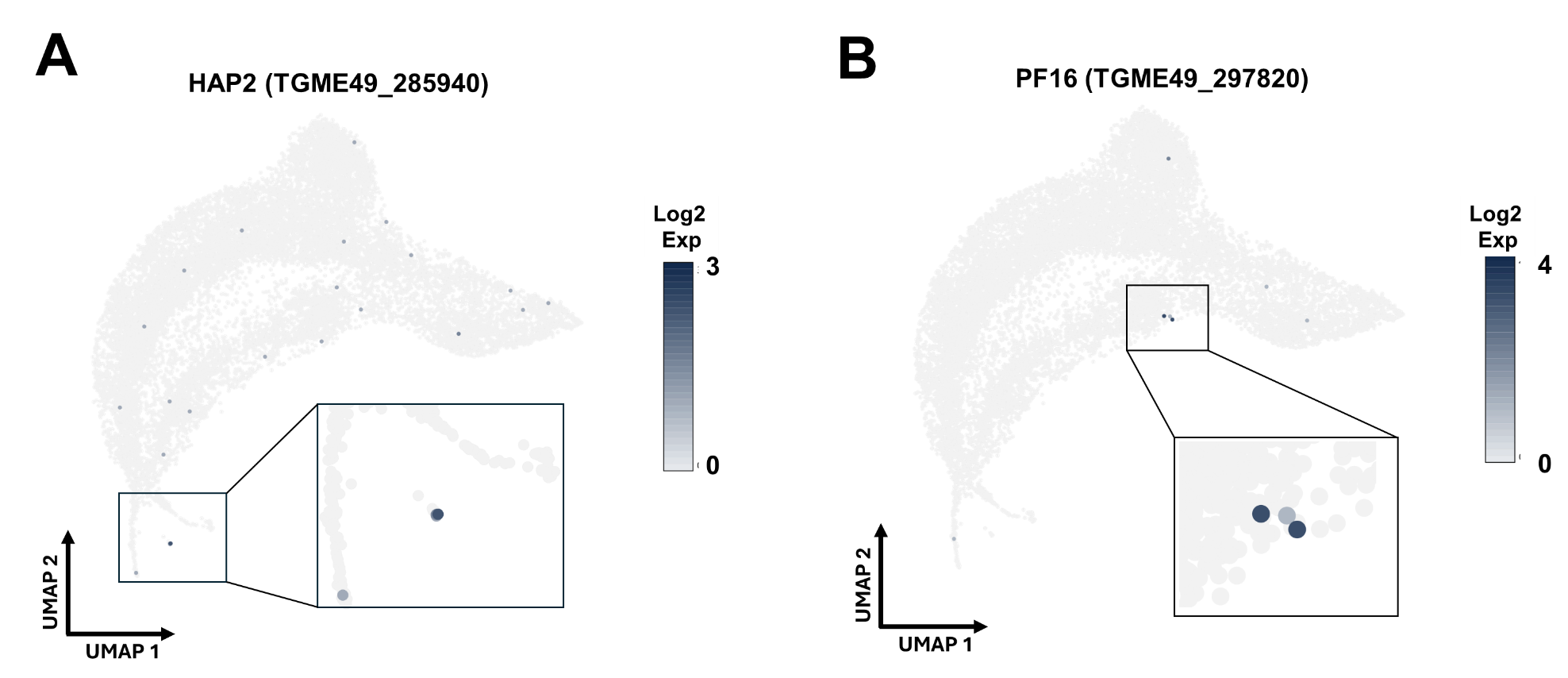

**Extended Data Figure 5:** **A)** Feature plot showing transcript abundance for the *HAP2* gene **(A)** or the *PF16* gene, two previously identified male-specific transcripts. Transcripts for HAP2 are detected at highest abundance in the “Male-1” cluster (original cluster 8) while PF16 is uniquely expressed in the “Male-2” cluster, a cluster of 3 cells (originally called cluster 9) with multiple male-specific transcripts.

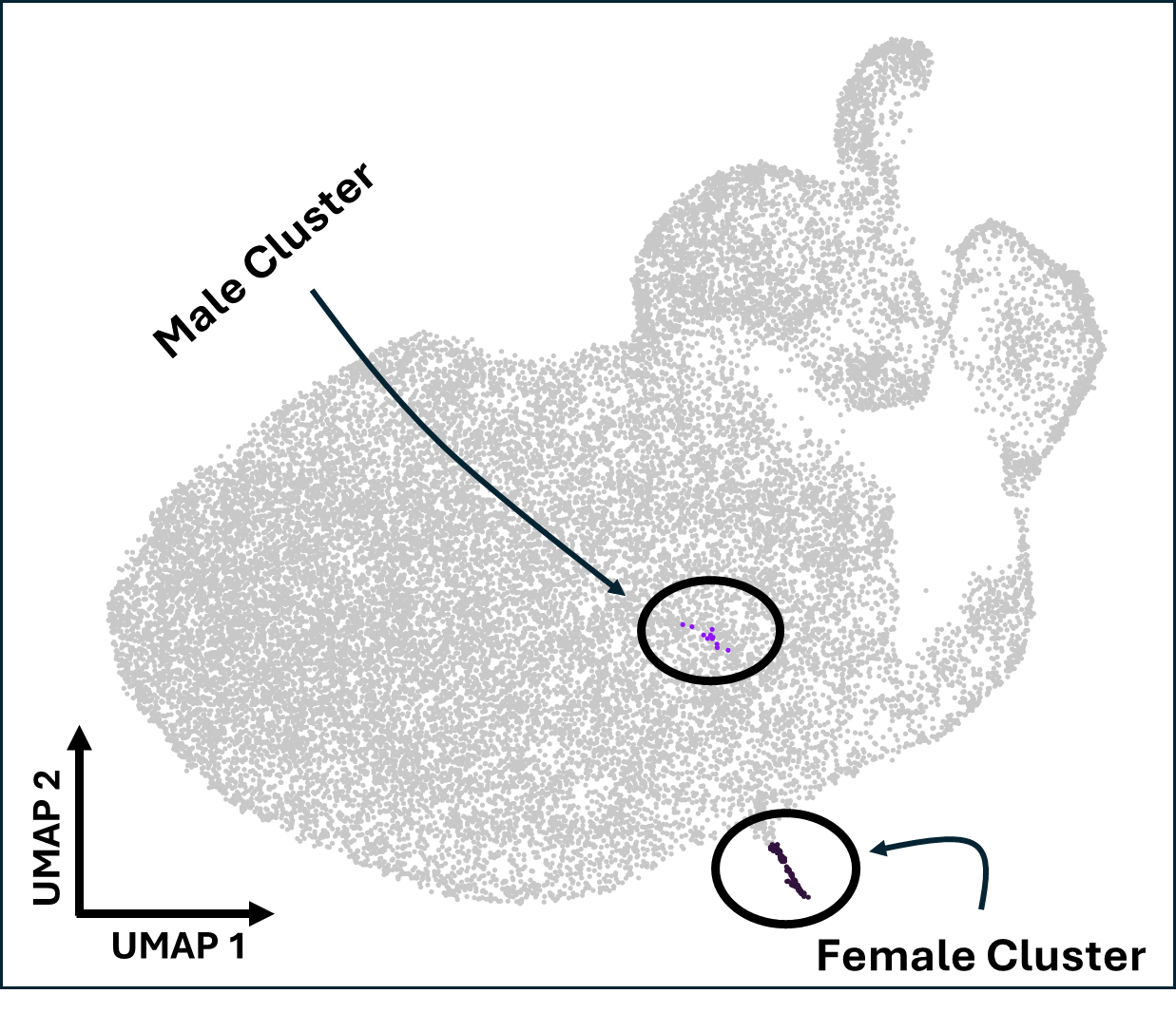

**Extended Data Figure 6:** Analysis of scRNAseq data derived from cells harvested at 7 DPI showing a marked imbalance between the numbers of male and female gametocytes, with female cells greatly outnumbering male cells (106 vs. 11). This observation, taken from cells isolated from the perturb-seq experiment, is consistent with data from cats 1 and 2 and with previous reports of sex ratio bias in Apicomplexan sexual stages. Cells were annotated based on transcript abundance for reported male and female gametocyte markers as described in the text.

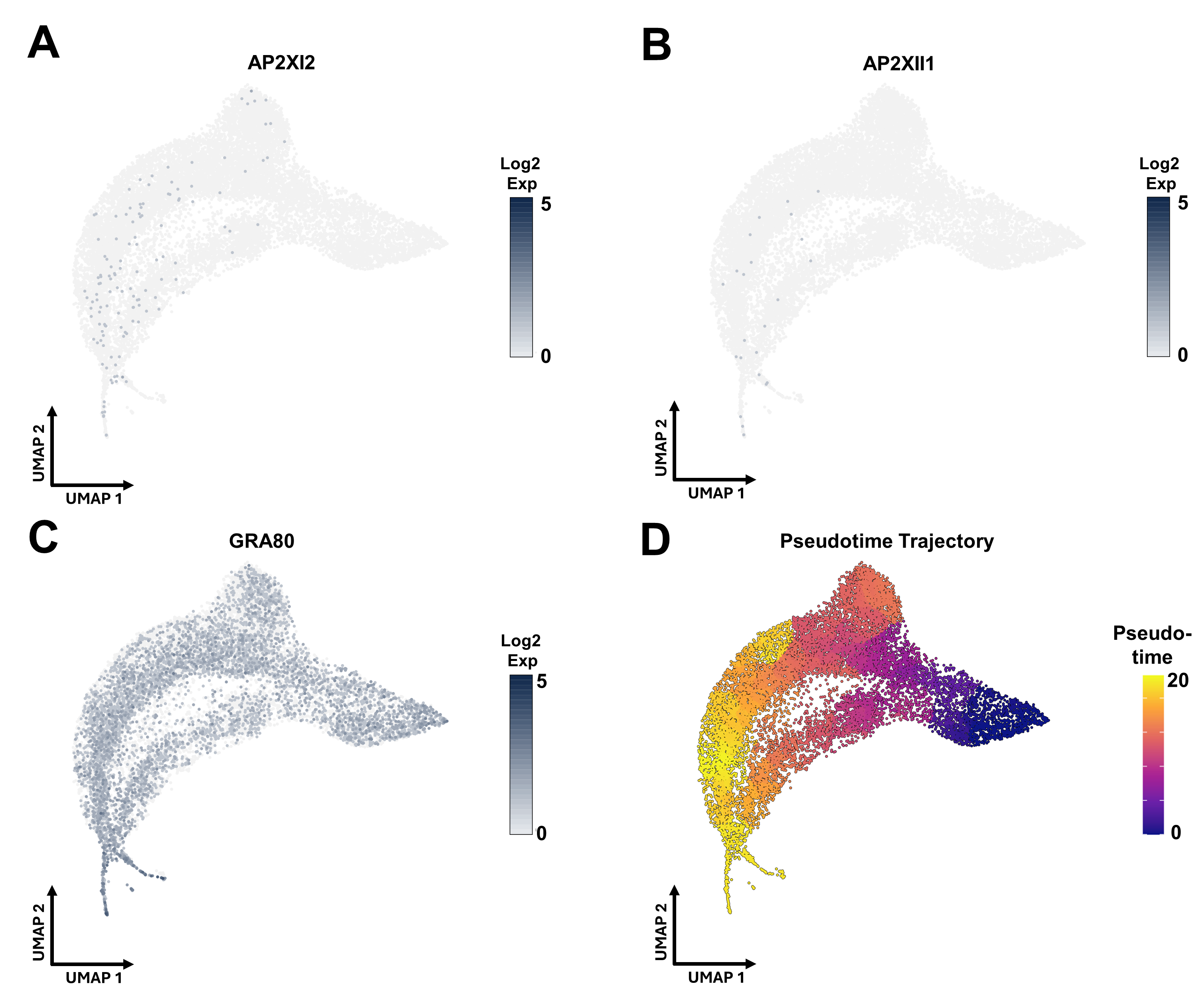

**Extended Data Figure 7:** **A-C)** Feature plots showing transcript abundance for AP2XI2 and AP2XII1 (**A** and **B**) which are both poorly expressed in a relatively small number of cells compared to a GRA80 **(C)**. **D)** Pseudotime analysis of scRNA-seq data using Monocle3. The inverse of gamete cells was used as the root for trajectory inference.

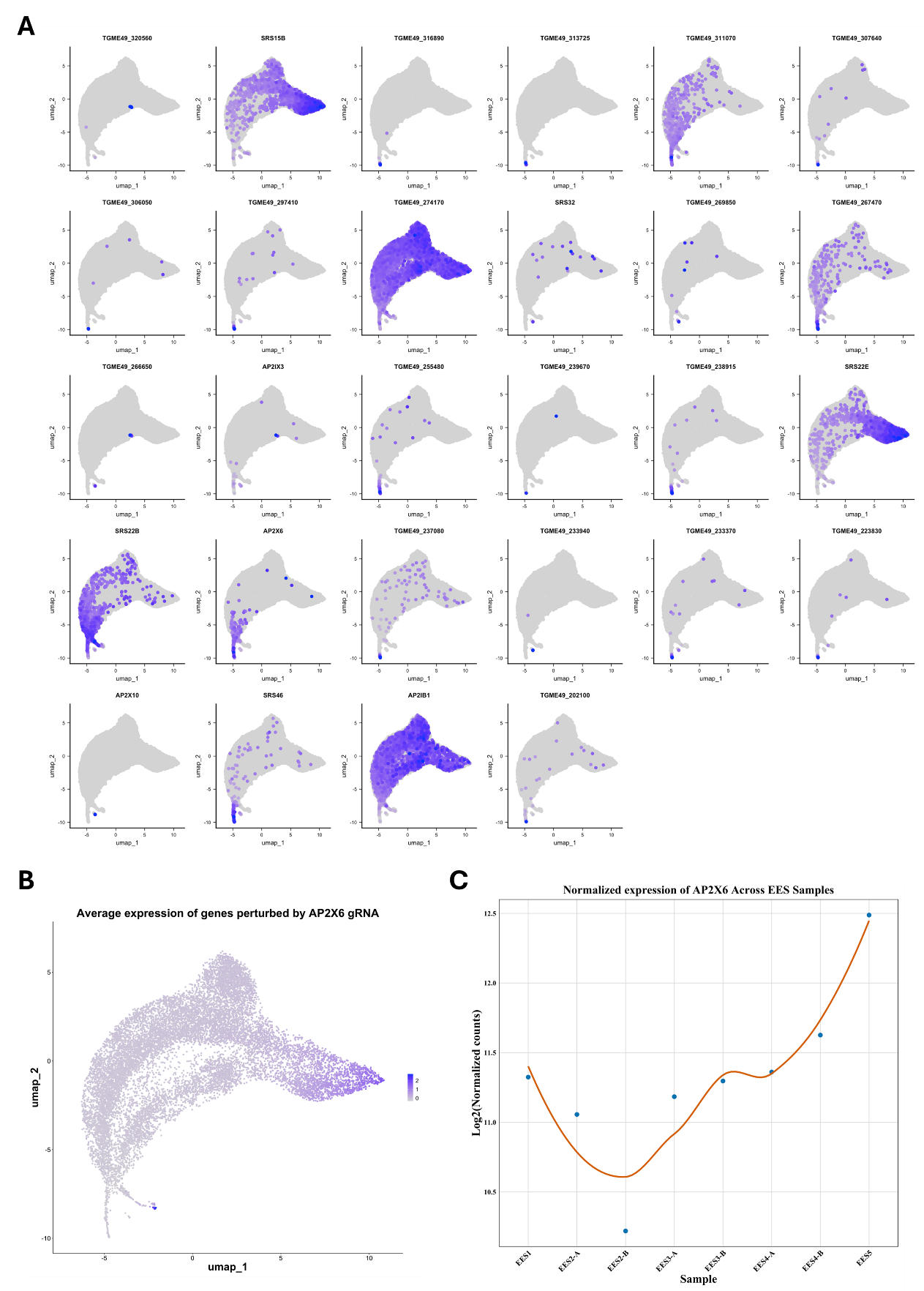

**Extended Data Figure 8:** **A)** Expression profiles for the 28 candidate genes included in the perturb-seq screen. B) Summed transcript abundance for the top 100 genes that were significantly altered by the presence of the gRNA targeting AP2X6. C) Previously published data from Ramakrishnan *et al.* (2019) [^1^](https://sciwheel.com/work/citation?ids=8641986&pre=&suf=&sa=0&dbf=0) showing that AP2X6 is of highest transcript abundance in later development clusters of cat enteroepithelial stages.

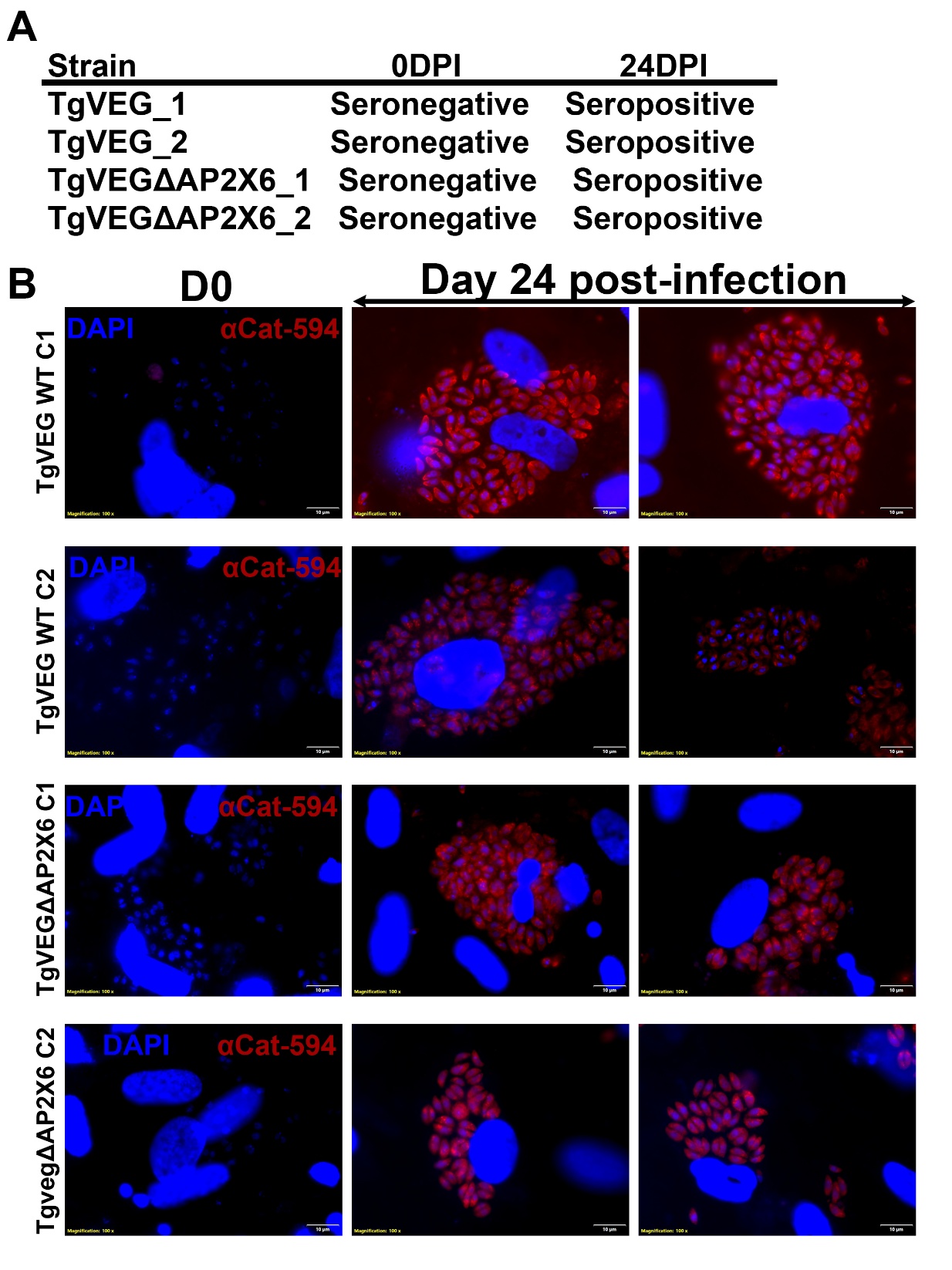

**Extended Data Figure 9: A)** Summary table of immunofluorescence results using cat serum taken pre (D0) and post (Day 24)-infection with WT or ΔAP2X6 *T. gondii* tissue cysts. **B)** Representative images for serum taken from each cat and used in immunofluorescence experiments at a 1:100 dilution followed by staining with goat anti-Cat antibody coupled to Alexa Fluor 594. Coverslips were pre-seeded with HFFs and infected with RH-YFP for 24 hours prior to fixation and staining. Dapi was used to stain parasite and host cell nuclei and only the blue and red channels are shown for clarity. One field of view is shown for D0 serum samples and two for D24 serum samples.
