## Supplementary Tables for "A single-cell atlas of *Toxoplasma* sexual development in the feline intestinal tract"

**Table 1.** 108 female gametocyte–specific genes shared among *T. gondii*, *C. parvum*, and *P. berghei*.

| | Gene ID |
|---|---|
| 1 | TGME49_201150 |
| 2 | TGME49_202880 |
| 3 | TGME49_204420 |
| 4 | TGME49_204560 |
| 5 | TGME49_205490 |
| 6 | TGME49_205658 |
| 7 | TGME49_206640 |
| 8 | TGME49_209470 |
| 9 | TGME49_210260 |
| 10 | TGME49_210950 |
| 11 | TGME49_211350 |
| 12 | TGME49_213340 |
| 13 | TGME49_214440 |
| 14 | TGME49_214580 |
| 15 | TGME49_215500 |
| 16 | TGME49_216380 |
| 17 | TGME49_216400 |
| 18 | TGME49_216890 |
| 19 | TGME49_218560 |
| 20 | TGME49_220570 |
| 21 | TGME49_221320 |
| 22 | TGME49_221350 |
| 23 | TGME49_221830 |
| 24 | TGME49_223150 |
| 25 | TGME49_223700 |
| 26 | TGME49_224060 |
| 27 | TGME49_225050 |
| 28 | TGME49_225220 |
| 29 | TGME49_226230 |
| 30 | TGME49_227430 |
| 31 | TGME49_227570 |
| 32 | TGME49_227580 |
| 33 | TGME49_228680 |
| 34 | TGME49_230420 |
| 35 | TGME49_230430 |
| 36 | TGME49_230590 |
| 37 | TGME49_231420 |
| 38 | TGME49_236070 |
| 39 | TGME49_238200 |
| 40 | TGME49_239890 |
| 41 | TGME49_240575 |
| 42 | TGME49_243600 |
| 43 | TGME49_244620 |

44 TGME49_244850
45 TGME49_245610
46 TGME49_247690
47 TGME49_248730
48 TGME49_248960
49 TGME49_249200
50 TGME49_250360
51 TGME49_250880
52 TGME49_253030
53 TGME49_253150
54 TGME49_253480
55 TGME49_254760
56 TGME49_256040
57 TGME49_258400
58 TGME49_259590
59 TGME49_261560
60 TGME49_262710
61 TGME49_263410
62 TGME49_264070
63 TGME49_267410
64 TGME49_267680
65 TGME49_268310
66 TGME49_269120
67 TGME49_269400
68 TGME49_271892
69 TGME49_272900
70 TGME49_273040
71 TGME49_273380
72 TGME49_273705
73 TGME49_276155
74 TGME49_278660
75 TGME49_279440
76 TGME49_282020
77 TGME49_285470
78 TGME49_285690
79 TGME49_285920
80 TGME49_286250
81 TGME49_287490
82 TGME49_288220
83 TGME49_289050
84 TGME49_290610
85 TGME49_290870
86 TGME49_290930
87 TGME49_291860
88 TGME49_294820

| | |
|---|---|
| 89 | TGME49_298630 |
| 90 | TGME49_301370 |
| 91 | TGME49_304750 |
| 92 | TGME49_307570 |
| 93 | TGME49_307580 |
| 94 | TGME49_310000 |
| 95 | TGME49_310660 |
| 96 | TGME49_313950 |
| 97 | TGME49_314500 |
| 98 | TGME49_316140 |
| 99 | TGME49_318540 |
| 100 | TGME49_319700 |
| 101 | TGME49_320500 |
| 102 | TGME49_326800 |
| 103 | TGME49_500391 |
| 104 | TGME49_286778 |
| 105 | TGME49_286782 |
| 106 | TGME49_313725 |
| 107 | TGME49_311070 |
| 108 | TGME49_239670 |

**Table 2.** Putative microgamete marker genes reported by Ramakrishnan *et al*. Scientific Reports 9, 1474 (2019)

| | Gene ID |
|---|---|
| 1 | TGME49_239550 |
| 2 | TGME49_258880 |
| 3 | TGME49_291160 |
| 4 | TGME49_201750 |
| 5 | TGME49_254020 |
| 6 | TGME49_261022 |
| 7 | TGME49_203135 |
| 8 | TGME49_243482 |
| 9 | TGME49_249840 |
| 10 | TGME49_273478 |
| 11 | TGME49_254180 |
| 12 | TGME49_255200 |
| 13 | TGME49_233330 |
| 14 | TGME49_237490 |
| 15 | TGME49_260000 |
| 16 | TGME49_297820 |
| 17 | TGME49_283765 |
| 18 | TGME49_249365 |
| 19 | TGME49_253500 |
| 20 | TGME49_250720 |
| 21 | TGME49_293890 |
| 22 | IFT81 |
| 23 | TGME49_215180 |
| 24 | TGME49_315845 |
| 25 | TGME49_270210 |
| 26 | TGME49_286660 |
| 27 | TGME49_211910 |
| 28 | TGME49_285410 |
| 29 | TGME49_230135 |
| 30 | TGME49_310480 |
| 31 | TGME49_212075 |
| 32 | TGME49_243550 |
| 33 | TGME49_293850 |
| 34 | TGME49_297670 |
| 35 | TGME49_217390 |
| 36 | TGME49_286932 |
| 37 | TGME49_230830 |
| 38 | TGME49_272360 |
| 39 | TGME49_217650 |
| 40 | TGME49_248225 |
| 41 | TGME49_207410 |
| 42 | TGME49_273472 |
| 43 | TGME49_312980 |

44 TGME49_233940
45 TGME49_202970

**Table 3.** List of primers used in this study

| **Primers** | **Sequence** |
|---|---|
| GRA11_promoter_fwd | accagtactacagccttcgagaaatgatttctttcgcaag |
| GRA11_promoter_rev2 | cgcccttggacattttgactttgtcaacgaac |
| mCherry_fwd2 | caaagtcaaaatgtccaagggcgaagaggac |
| mCherry_rev | cgggcagcttctgtactgcactacttgtacagctcgtccattc |
| Pclick_uprt_R-homology_arm | CCTTcTATTCCAAGATCTGTCTCGAGGTCGACTGGAACTAC |
| Pclick_uprt_L-homology_arm | cgCTTTCCATCGACTCGCCGCCGCTCTAGAACTAGTGGAT |
| Capture-Sequence1F | ggtcctagcaaggccAAGTGGCACCGAGTC |
| Capture-Sequence1R | ggccttaaagcggccAAGTTGATAACGGACTAGC |

**Table 4.** gRNAs used in this study.

| Target | Name | Sequence |
|---|---|---|
| TGME49_320560 | p25alphafamilyprote_TGME49_320560-1 | GGCGGCTATCGATCTCATCG |
| TGME49_320560 | p25alphafamilyprote_TGME49_320560-2 | CCAGTGGTCCGGTGTATAGT |
| TGME49_320560 | p25alphafamilyprote_TGME49_320560-3 | GGCGGTCCGAGCACCGTTGA |
| SRS15B | SRS15B_TGME49_320240-1 | TTTCCGTGTGCACCTGATGC |
| SRS15B | SRS15B_TGME49_320240-2 | CAAATGGTAGCTACGGTCTC |
| SRS15B | SRS15B_TGME49_320240-3 | GGTAGCTACGGTCTCTGGTG |
| TGME49_316890 | hypprot_TGME49_316890-1 | TAGGTCCACGGAGACATCGT |
| TGME49_316890 | hypprot_TGME49_316890-2 | TCACGTTGGTACTGGCTTCG |
| TGME49_316890 | hypprot_TGME49_316890-3 | GCACAGAACGCATACGGGAG |
| TGME49_313725 | hypprot_TGME49_313725-1 | ACAGGAACGACAAAGGCTGT |
| TGME49_313725 | hypprot_TGME49_313725-2 | GCACTGCGACTGGTTGTGAC |
| TGME49_313725 | hypprot_TGME49_313725-3 | CTGCTTACACTAGCGACCAG |
| TGME49_311070 | hypprot_TGME49_311070-1 | ATCGGGGCGGGGCAGCGCAA |
| TGME49_311070 | hypprot_TGME49_311070-2 | AGACTCAGAGGACTTCACTG |
| TGME49_311070 | hypprot_TGME49_311070-3 | CGTCTGGAAAGGGTGCCCCG |
| TGME49_307640 | CMGCkinase2CCK2fami_TGME49_307640-1 | CAACTTCTTCGGGAGTCATT |
| TGME49_307640 | CMGCkinase2CCK2fami_TGME49_307640-2 | GTGAGGAACGCGTAGGTACG |
| TGME49_307640 | CMGCkinase2CCK2fami_TGME49_307640-3 | CTGATCCAGGAAGACGTTTG |
| TGME49_306050 | hypprot_TGME49_306050-1 | ATTGGCTGCGGCATTCTGTG |
| TGME49_306050 | hypprot_TGME49_306050-2 | TCTGTTTGAGGGAAGCACAA |
| TGME49_306050 | hypprot_TGME49_306050-3 | TTTCCCGCCTTGATTTCCTC |
| TGME49_297410 | hypprot_TGME49_297410-1 | GAAGTATGTCCTGATGTTGG |
| TGME49_297410 | hypprot_TGME49_297410-2 | TATCGACCTCCAGCCGAAAC |
| TGME49_297410 | hypprot_TGME49_297410-3 | CCTGCCTATTAGTTCTCTCG |
| TGME49_274170 | proteinkinaseincomp_TGME49_274170-1 | AAACTCTGGACGCCCTACTC |
| TGME49_274170 | proteinkinaseincomp_TGME49_274170-2 | CATCGCAGCGAGCTTTGCCA |
| TGME49_274170 | proteinkinaseincomp_TGME49_274170-3 | TCGTTGGGTGGAGATTGCCG |
| SRS32 | SRS32_TGME49_271980-1 | GGTGTTTCCACATGTCATCG |
| SRS32 | SRS32_TGME49_271980-2 | ATACCTCGACGGGCCAGTTT |
| SRS32 | SRS32_TGME49_271980-3 | CAAGTCCCGCCGCGCACCTG |
| TGME49_269850 | proteinc21orf592Cpu_TGME49_269850-1 | TGTCTGCCCACGGGCCCTTG |
| TGME49_269850 | proteinc21orf592Cpu_TGME49_269850-2 | TGTTTCCTCCAGAATTTCAG |
| TGME49_269850 | proteinc21orf592Cpu_TGME49_269850-3 | CTTCTCGTGCGAAACAGTCT |
| TGME49_267470 | coldshockDNAbinding_TGME49_267470-1 | GCGAGATGGCAGTGTCACGC |
| TGME49_267470 | coldshockDNAbinding_TGME49_267470-2 | GCTACCAGCAACATTTGCGT |
| TGME49_266650 | hypprot_TGME49_266650-1 | AGTTAAGGCAGAGATTCCAG |
| TGME49_266650 | hypprot_TGME49_266650-2 | AAACGTGACACCTCTCACGG |
| AP2IX3 | AP2IX3_TGME49_264485-1 | TCCCGGCACGGAGTCTGTTG |
| AP2IX3 | AP2IX3_TGME49_264485-2 | AGAGTAAAAGGTCTCGATTG |
| AP2IX3 | AP2IX3_TGME49_264485-3 | TTGCGGCAGCGAGACCGCAG |
| TGME49_255480 | thioredoxindomainco_TGME49_255480-1 | CTCCAGGTCAGACTGCAACG |
| TGME49_255480 | thioredoxindomainco_TGME49_255480-2 | AATGGACCATGCGCGTAACG |
| TGME49_255480 | thioredoxindomainco_TGME49_255480-3 | ATTCTGAGATGGGAGTTAGC |

| | | |
|---|---|---|
| TGME49_239670 | hypprot_TGME49_239670-1 | CAAGTCGCGCAGGATGAAAG |
| TGME49_239670 | hypprot_TGME49_239670-2 | AAAGTCAACACGATCCTGCC |
| TGME49_239670 | hypprot_TGME49_239670-3 | AGCAGCTCGTGGAACCCTTG |
| TGME49_238915 | hypprot_TGME49_238915-1 | AGACTGTTGAGATTCAGCAC |
| TGME49_238915 | hypprot_TGME49_238915-2 | TTTGAGACTGCATCTCCAAG |
| TGME49_238915 | hypprot_TGME49_238915-3 | CCACTGCAGAAATCTCCAAT |
| SRS22E | SRS22E_TGME49_238490-1 | CGTCTTCACCGCAAGTGAAG |
| SRS22E | SRS22E_TGME49_238490-2 | GGAGTGCTGGCGTTGAACGC |
| SRS22E | SRS22E_TGME49_238490-3 | ACTAGAAAATGTCACGACGG |
| SRS22B | SRS22B_TGME49_238460-1 | TCGGTGATGTTGAAGCTGAG |
| SRS22B | SRS22B_TGME49_238460-2 | CTTCGGGTCCAGGTTCTTCA |
| SRS22B | SRS22B_TGME49_238460-3 | GTTAACCTGGCAATCGTTGG |
| AP2X6 | AP2X6_TGME49_237425-1 | TCAGGGGCCCCGTCGCCGAG |
| AP2X6 | AP2X6_TGME49_237425-2 | GTTCTCATACCGACTGTCTG |
| AP2X6 | AP2X6_TGME49_237425-3 | TTAGGTGTCTTCTTTCCACG |
| TGME49_237080 | hypprot_TGME49_237080-1 | TGTCTCAGTTTCAGTCGCAG |
| TGME49_237080 | hypprot_TGME49_237080-2 | CTTGGGAAAGAAGAGCCGTG |
| TGME49_237080 | hypprot_TGME49_237080-3 | GGCCTTCTTCTGAACAGGCG |
| TGME49_233940 | leucinerichrepeatco_TGME49_233940-1 | TCAGTTGTTGGAGTCCCTGT |
| TGME49_233940 | leucinerichrepeatco_TGME49_233940-2 | GGAGTCCGGTGAATTCTTCA |
| TGME49_233940 | leucinerichrepeatco_TGME49_233940-3 | CGAGGTACGTCAATTCTTTC |
| TGME49_233370 | hypprot_TGME49_233370-1 | TGAGTTGCATGGCCTTCCAA |
| TGME49_233370 | hypprot_TGME49_233370-2 | TTAATTCTCCATCTGGTCCG |
| TGME49_233370 | hypprot_TGME49_233370-3 | CTCCGTCCATAAGCGCCTGG |
| TGME49_223830 | fasciclindomaincont_TGME49_223830-1 | CGTGGTATCCACCTCCATTG |
| TGME49_223830 | fasciclindomaincont_TGME49_223830-2 | AATGGAGGTGGATACCACGT |
| TGME49_223830 | fasciclindomaincont_TGME49_223830-3 | TGTCTGATCTAACTCCTCCT |
| AP2X10 | AP2X10_TGME49_215340-1 | AGTAGCAGCGGTATCTTTAG |
| AP2X10 | AP2X10_TGME49_215340-2 | CAACGACAGCAACAACTTCG |
| AP2X10 | AP2X10_TGME49_215340-3 | AAGTTGTTGCTGTCGTTGTG |
| SRS46 | SRS46_TGME49_214190-1 | TCGCGGCTCCGAGTTTCGCG |
| SRS46 | SRS46_TGME49_214190-2 | GACTGTCGCGAAGATGGTCG |
| SRS46 | SRS46_TGME49_214190-3 | TTGGGAGCCAGGCAAAGCTG |
| AP2IB1 | AP2IB1_TGME49_208020-1 | CGCCTCCCGCCAATCGTCCG |
| AP2IB2 | AP2IB1_TGME49_208020-2 | GTTCGACAAATGTGCACCTG |
| AP2IB3 | AP2IB1_TGME49_208020-3 | CTTGCCCTCAGAACTCGTTG |
| TGME49_202100 | hypprot_TGME49_202100-1 | TGATGGATCGCAGTGTGTCT |
| TGME49_202100 | hypprot_TGME49_202100-2 | ATTGGAGGCGGCTCCAGTTT |
| TGME49_202100 | hypprot_TGME49_202100-3 | CGCCTCCAATGCCTCCAATG |
| Non-Targeting | NEG_CTRL-1 | GGATTGAATGGCTAACGCGG |
| Non-Targeting | NEG_CTRL-2 | TGCACCCACCGTAGTTATCG |
| Non-Targeting | NEG_CTRL-3 | GTATTACTGATATTGGTGGG |
